# Abnormal social behaviors and dysfunction of autism-related genes associated with daily agonistic interactions in mice

**DOI:** 10.1101/125674

**Authors:** Natalia N. Kudryavtseva, Irina L. Kovalenko, Dmitry A. Smagin, Anna G. Galyamina, Vladimir N. Babenko

## Abstract

**Background:** The ability of people to communicate with each other is a necessary component of social behavior and the normal development of individuals who live in a community. An apparent decline in sociability may be the result of a negative social environment or the development of affective and neurological disorders, including autistic spectrum disorders. The behavior of these humans may be characterized by the deterioration of socialization, low communication, and repetitive and restricted behaviors. This study aimed to analyze changes in the social behaviors of male mice induced by daily agonistic interactions and investigate the involvement of genes, related with autistic spectrum disorders in the process of the impairment of social behaviors.

**Methods:** Abnormal social behavior is induced by repeated experiences of aggression accompanied by wins (winners) or chronic social defeats (losers) in daily agonistic interactions in male mice. The collected brain regions (the midbrain raphe nuclei, ventral tegmental area, striatum, hippocampus, and hypothalamus) were sequenced at JSC Genoanalytica (http://genoanalytica.ru/, Moscow, Russia). The Cufflinks program was used to estimate the gene expression levels. Bioinformatic methods were used for the analysis of differentially expressed genes in male mice.

**Results:** The losers exhibited an avoidance of social contacts toward unfamiliar conspecific, immobility and low communication on neutral territory. The winners demonstrated aggression and hyperactivity in this condition. The exploratory activity (rearing) and approaching behavior time towards the partner were decreased, and the number of episodes of repetitive self-grooming behavior was increased in both social groups. These symptoms were similar to the symptoms observed in animal models of autistic spectrum disorders. In an analysis of the RNA-Seq database of the whole transcriptome in the brain regions of the winners and losers, we identified changes in the expression of the following genes, which are associated with autism in humans: *Tph2, Maoa, Slc6a4, Htr7,Gabrb3, Nrxn1, Nrxn2, Nlgn1, Nlgn2, Nlgn3, Shank2, Shank3, Fmr1, Ube3a, Pten, Cntn3, Foxp2, Oxtr, Reln, Cadps2, Pcdh10, Ctnnd2, En2, Arx, Auts2, Mecp2*, and *Ptchd1*.Common and specific changes in the expression of these genes in different brain regions were identified in the winners and losers.

**Conclusions:** This research demonstrates for the first time that abnormalities in social behaviors that develop under a negative social environment in adults may be associated with alterations in expression of genes, related with autism in the brain.

## Introduction

The ability of people to communicate with each other is a necessary component of social behavior and the normal development of individuals who live in a community. An apparent decline in sociability may be the result of a negative social environment or the development of affective and neurological disorders. A pronounced impairment in communication is associated with autism, which is regarded as a neural development disorder with an onset at an early age [1]. Autism is characterized by impaired socialization, low communication and restricted and/or repetitive behaviors [2], which are accompanied by many other neurobiological symptoms [3]. Moreover, there is a growing consensus that autism comprises a complex syndrome and several causes act simultaneously [1], which lead to the autism spectrum disorders (ASDs) that have roots in the early development stages [4, 5] and arise, for example, under prenatal stress [6]. Autistic risks are thought to include an unbalanced diet, specific diseases, toxic exposure, infectious diseases, stress and a multitude of other causes [7–8]. In the latter case, low communication may persist and acquire a chronic course. Furthermore, the symptoms of ASDs may accompany the development of various diseases [2]. There is comorbidity of ASDs with several disorders, such as epilepsy, schizophrenia, ADHD, aggression, depression, and anxiety [9–12].

To date, experimental studies have been focused on the identification of adequate models to investigate the mechanisms and corrections of impaired social behaviors. Animal studies have demonstrated that social impairment in rodents may be produced by prenatal stress [13, 14], early handling [15], maternal separation [16], and early social isolation [17, 18]. Several knockouts or mutant mice, such as 4E-BP2, Fmr1, Mecp2, Ube3a, Nf1, Pten, Tsc1/Tsc2, and Shank, exhibited low sociability compared with wild type animals [19-23]. Among the strains of animals that exhibit low communication, the BTBR T+tf/J, BALB/cJ, C57BL/6J, A/J and 129S1/SvImJ mouse lines have been considered [24-28]. Thus, genetic influence and/or social environment may lead to the impairment of social behaviors.

The communicative ability of animals is typically evaluated at the time of behavioral response to another individual in different experimental situations in the home cage or neutral territories [25, 27, 29]. If animals of one strain exhibited a reduced response to the tester compared with animals of other strains, these animals were considered suitable for investigating the mechanisms of decreased social behavior and its treatments. Knockouts and wild type controls were also compared. Other changes in social behaviors and repetitive behaviors [25, 26, 28] were also considered in animals. In animal studies, attention is also focused on reduced activity or low sociability, the deterioration of social recognition, and increased anxiety. Restricted (repetitive) behaviors, such as increased motor activities (spontaneous activity, exploration, circling, digging, and jumping) and increased self-involved behaviors (self-grooming, scratching, and circling behavior) [25, 26, 28] are also considered symptoms of social behavior impairments in animals.

Our studies have previously demonstrated that male mice with long-term, daily, repeated experiences of aggression or chronic social defeats exhibited disturbances in social recognitions and motivated behaviors, pronounced anxiety and aggressiveness, a depression-like state, and stereotypic behaviors [30–36]. The initial findings indicated that these mice – winners and losers – demonstrate disturbances in communication and sociability [5, 32, 37]. This study aimed to analyze in detail the social behaviors of male mice that chronically live in a hostile environment with the goal to identify associations with forms of behavior that other authors [25, 26, 28] have considered as autistic spectrum symptoms similar to behaviors in humans.

The hypothesis was that autistic symptoms may develop in adult mice in a hostile environment in our behavioral paradigm. We also suggested that changes in social behaviors may involve changes in the expression of genes associated with autism in the brain. To investigate this hypothesis, we used our RNA-Seq analysis of the whole transcriptome in the midbrain raphe nuclei, ventral tegmental area (VTA), striatum, hippocampus, and hypothalamus of the winners and losers to identify changes in the expression of recognized autism-related genes that have strong associations, according to databases (OMIM, http://omim.org/; MalaCards, http://www.malacards.org), with autism or autistic spectrum disorders, namely, *Tph2, Maoa, Slc6a4, Htr7, Shank2* (*auts17*), *Shank3, Mecp2* (*autsx3*), *Ube3a, Gabrb3, Cntn3, Reln, Pten, Ctnnd2, Nrxn1, Nrxn2, Nlgn1, Nlgn2, Nlgn3* (*autsx1*), *Pcdh10, Cadps2, Foxp2, Fmr1, Auts2, En2, Arx, Oxtr*, and *Ptchd1*(*autsx4*). Most genes of interest are related to the signal neurotransmission network.

## Materials and Methods

### Animals

Adult male mice of the C57BL/6J strain from the Animal Facility of the Institute of Cytology and Genetics, SD RAS, (Novosibirsk, Russia) were used. The animals were housed under standard conditions (12:12 h light/dark regime, switch-on at 8.00 a.m.; food (pellets) and water available *ad libitum*). Mice were weaned at one month of age and housed in groups of 8-10 in plastic cages (36 × 23 × 12 cm). Experiments were performed on mice 10-12 weeks of age. All procedures were in compliance with the European Communities Council Directive 210/63/EU on September 22, 2010. The study was approved by Scientific Council N 9 of the Institute of Cytology and Genetics SD RAS of March, 24, 2010, N 613.

### Generation of positive and negative social experience in male mice

Prolonged positive and negative social experience (wins and defeats) in male mice was induced by daily agonistic interactions [38, 39]. Pairs of weight-matched animals were each placed in a steel cage (14 × 28 × 10 cm) bisected by a perforated transparent partition allowing the animals to see, hear and smell each other, but preventing physical contact. The animals were left undisturbed for two or three days to adapt to new housing conditions and sensory contact before they were exposed to encounters. Every afternoon (14:00-17:00 p.m. local time), the cage lid was replaced by a transparent one, and 5 min later (the period necessary for individuals’ activation), the partition was removed for 10 minutes to encourage agonistic interactions. The superiority of one of the mice was firmly established within two or three encounters with the same opponent. The superior mouse would be attacking, biting and chasing another, who would be displaying only defensive behavior (sideways postures, upright postures, withdrawal, lying on the back or freezing). As a rule, aggressive interaction between males are discontinued by lowering the partition if the sustained attacks has lasted 3 min (in some cases less) thereby preventing the damage of losers. Each defeated mouse (defeater, loser) was exposed to the same winner for three days, while afterwards each loser was placed, once a day after the fight, in an unfamiliar cage with an unfamiliar winner behind the partition. Each victorious mouse (winners, aggressors) remained in its original cage. This procedure was performed once a day for 20 days and yielded an equal number of winners and losers.

In the behavioral study, the following three groups of animals were used: (1) Losers – groups of chronically defeated mice after 10 and 20 days of agonistic interactions (Los-10 and Los-20); (2) Winners – groups of chronically victorious mice that exhibit daily aggression during 10 and 20 days (Win-10 and Win-20) of agonistic interactions; (3). Controls – mice without a consecutive experience of agonistic interactions [38, 39].

Twenty-day winners and 20-day losers with the most expressed behavioral phenotypes were selected for the transcriptome analysis. Different mice were used for the behavioral study and transcriptomic analysis. All mice were simultaneously decapitated, including 20- time winners and losers, 24 hours after the last agonistic interaction and the control animals. The brain regions were dissected by one experimenter according to the map presented in the Allen Mouse Brain Atlas [http://mouse.brain-map.org/static/atlas]. All biological samples were placed in RNAlater solution (Life Technologies, USA) and were stored at -70° C until sequencing.

The brain regions were selected for the analysis based on their functions and localization of neurons of neurotransmitter systems. These areas are as follows: the midbrain raphe nuclei, which comprises a multifunctional brain region and contains the majority of serotonergic neuronal pericaryons; the ventral tegmental area (VTA), which contains the bodies of dopaminergic neurons, is widely implicated in the natural reward circuitry of the brain and is important in cognition, motivation, drug addiction, and emotions related to several psychiatric disorders; the striatum, which is responsible for the regulation of motor activity and stereotypical behaviors and is also potentially involved in various processes; the hippocampus, which belongs to the limbic system, is essential for memory consolidation and storage, and plays important roles in neurogenesis and emotional mechanisms; and the hypothalamus, which regulates the stress reaction and many other physiological functions.

## Behavioral study

### Partition test

The partition test can be utilized in these studies as a tool for estimating behavioral reactivity of mice to a conspecific behind the transparent perforated partition dividing the experimental cage into equal parts [29]. The number of approaches to the partition, and the total time spent near it (moving near the partition, smelling and touching it with one or two paws, clutching and hanging, putting noses into the holes or gnawing the holes) were scored during 5 min as indices of reacting to the partner in the neighboring compartment of home cage. The time the males showed sideways position or “turning away” near the partition is not included in the total time.

The experimental procedure was as follows. Two mice (a winner and a loser) were for almost one day together in a home cage with a partition (familiar partner). Then, the steel cover of the cage was replaced by a transparent one. The behaviors were videotaped during 5 min.

### Social interactions test

The social interactions test was developed to quantitatively assess social behavior in animals versus a standard tester, i.e., a group-housed unfamiliar partner of the same strain, weight and age. Every tester was used one time. The tested mouse and standard tester were simultaneously placed in a neutral territory, in a clean cage 36×23×12 cm in size, in opposite corners of the cage. The cage did not have litter and fodder, which may distract the mouse from the partner.

During a 10-minute test, the following behavioral domains were analyzed in the animals: 1) *Avoidance behavior* – avoidance of the approaching partner, flight, or freeze at his approach. In this case, the male may freeze in the upright posture; 2) *Approach behavior* directed toward the unfamiliar partner: communication – a behavior that aims to get closer to the partner by approaching, sniffing at and following him; 3) *Attacks* (attacking, biting and chasing) of the partner; 4) *Aggressive grooming* (the tested male mounts onto the unfamiliar partner’s back, holds it down and spends substantial time licking and nibbling at the scruff of the neck; the partner is wholly immobilized or may stretch out the neck and freeze; *5*) *Rearing*, which is considered an exploratory activity; 6) *Immobility* and waiting, in which the animal sits in the corner or near the wall and watches for the partner; 7) *Locomotor activity* of animals in the cage without attention to the partner; 8) *Self-grooming* (body care activities, such as fur licking and head or body washing), which is regarded as an indicator of displacement activity in new circumstances. The total time and number of events, as well as the % of animals that demonstrated forms of behaviors during the test were measured. If an animal did not exhibit behaviors, all counts were recorded as zero. After each male, the cage was washed several times with water and was dried with a clean tissue paper.

In our experiments, in general, an activation period was used prior to the behavior testing. The animals were taken to a test room and remained there for 5 minutes so that they could adapt to the lighting conditions and behavioral activation. The animal behavior during the test was recorded, and the videos were processed.

### RNA-Seq

The collected samples were sequenced at JSC Genoanalytica (www.genoanalytica.ru, Moscow, Russia), and the mRNA was extracted using a Dynabeads mRNA Purification Kit (Ambion, Thermo Fisher Scientific, Waltham, MA, USA). cDNA libraries were constructed using the NEBNext mRNA Library PrepReagent Set for Illumina (New England Biolabs, Ipswich, MA USA) following the manufacturer’s protocol and were subjected to Illumina sequencing. More than 20 million reads were obtained for each sample. The resulting “fastq” format files were used to align all reads to the GRCm38.p3 reference genome using the TopHat aligner [40]. DAVID Bioinformatics Resources 6.7 (http://david.abcc.ncifcrf.gov) was used for the description of differentially expressed gene ontology. The Cufflinks program was used to estimate the gene expression levels in FPKM units (fragments per kilobase of transcript per million mapped reads) and subsequently identify the differentially expressed genes in the analyzed and control groups. Each brain area was considered separately for 3 vs 3 animals. Only annotated gene sequences were used in the following analysis. Genes were considered differentially expressed at *P* ≤ 0.05. q < 0.05 was also taken into consideration.

We have previously conducted studies of gene expression in males in similar experiments using the RT-PCR method with larger samples for each compared experimental group, i.e., winners and losers (> 10 animals). The direction and extent of changes in expressions of the *Tph2, Slc6a4, Bdnf, Creb1*, and *Gapdh* genes in the midbrain raphe nuclei of males compared with the control produced by two methods, including RT-PCR [41, 42] and RNA-Seq, are generally consistent. This finding suggests that the transcriptome analyses of the data provided by the Company Genoanalitika (http://genoanalytica.ru, Moscow) have been verified, and this method reflects the actual processes that occur in the brain under our experimental paradigm.

The Human Gene Database (http://www.genecards.org/); GeneMania database (http://www.genemania.org), Online Mendelian Inheritance in Man database (OMIM, http://omim.org/); Human Disease Database (MalaCards, http://www.malacards.org); and Human Biological Pathway Unification Database (http://pathcards.genecards.org/) were used for the description and analysis of the data obtained.

### Selection of target genes

In an analysis of the whole transcriptome using RNA-Seq in five brain regions of male mice with chronic social defeats or wins in daily agonistic interactions, we identified changes in the expression of numerous genes, which are considered candidates for autism according to the previously described databases. Notably, a substantial portion of the candidate genes corresponds to the neurospecific genes involved in the transmission of nerve impulses, in particular synapses with specialized genes. This report is focused on the analysis of genes, which are strongly associated the development of autism or ASDs in humans and are also highly neurospecific genes as follows: Serotonergic genes: *Tph2, Maoa, Slc6a4*, and *Htr7*; Genes of cell adhesions and synaptic proteins: *Reln, Pten, Ctnnd2, Nrxn1, Nrxn2, Nlgn1, Nlgn2, Nlgn3* (*autsx1*), *Shank2* (*auts17*) and *Shank3;* Genes that code receptors: *Gabrb3, Htr7, Oxtr*, and *Pcdh10*; Genes of brain development: *Mecp2* (*autsx3*), *Arx, Auts2, En2, Foxp2, Fmr1*, and *Ptchd1* (*autsx4*) codes transmembrane protein; Transcription factors: *Foxp2*; Translation repressors: *Fmr1*; and Factor of protein degradation: *Ube3a* gene. We built the interaction network using a Strings association network resource (http://string-db.org; 43). It demonstrates interactions between proteins that are coded by most of our genes related to the signal neurotransmission networks (Fig. 1), which include serotonergic genes and genes of cell adhesions and synaptic proteins. The String database located all serotonergic genes closely related to each other and the *Gabrb3* gene (Fig. 1). A significant association has previously been demonstrated between *Gabrb3* and serotonergic genes in the regulation of schizophrenia symptoms, such as delusions, hallucinations, mania, depression, and negative symptoms [44, 45]. Furthermore, we analyzed the expression of the *Bdnf* gene coding the protein, which is involved in neural plasticity and, as a consequence, many psychiatric disorders, particularly autism and ASDs [46, 47].

**Figure 1.**
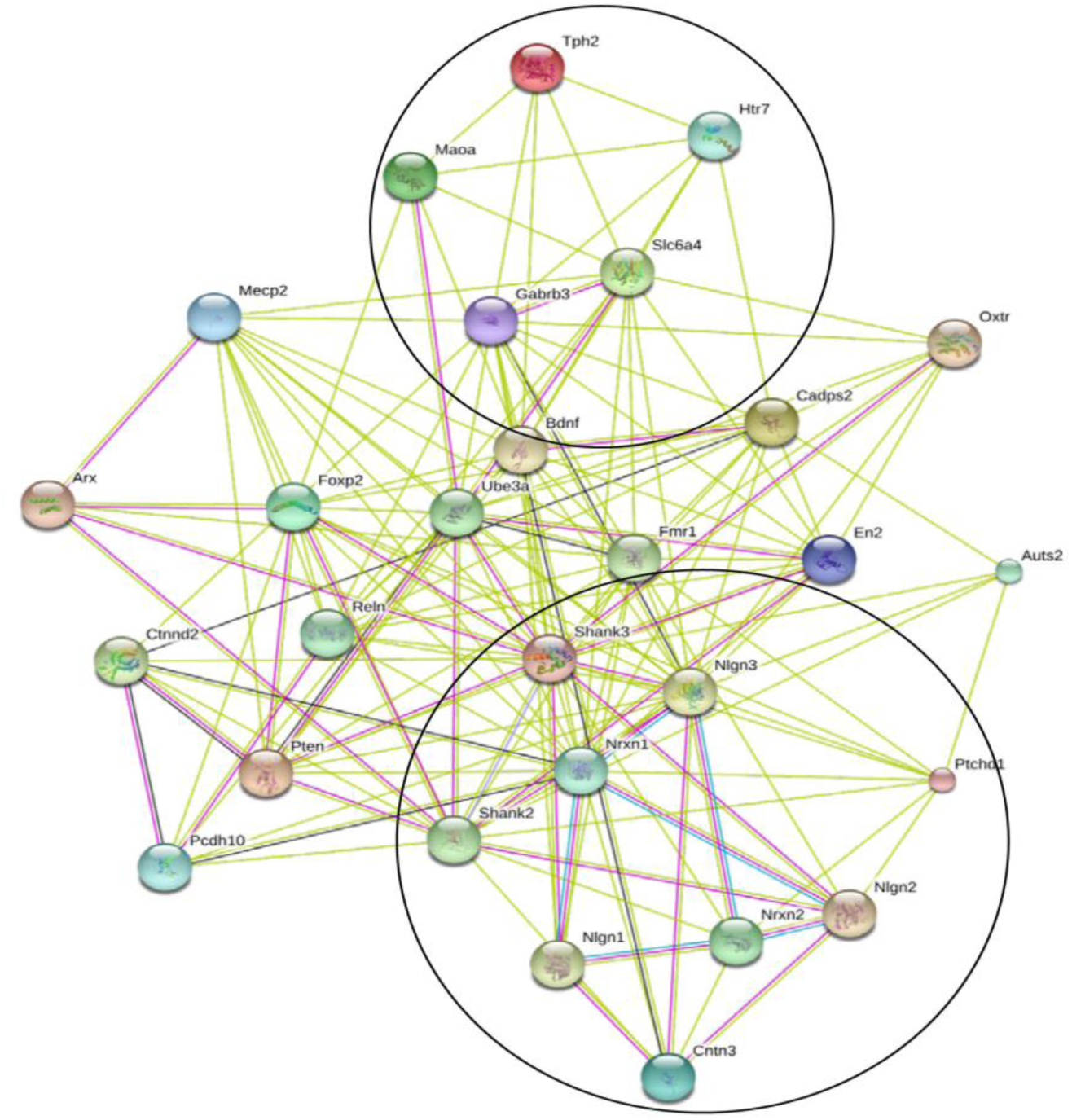
Gene association network based on the Strings text mining tool (http://string-db.org). Synaptic genes (low right) and the serotonergic (top center) system are encircled. All genes were connected on the first layer with no mediator second layer proteins. The majority of the associations is based on publication co-occurrence (salad color lines [43]); however, a significant number of the interactions is experimentally (red lines) confirmed. The center node corresponds to the *Shank3* gene, which maintains multiple experimentally confirmed interactions with other genes.

### Statistical analysis

Statistical analysis of the behavioral data was performed using one-way ANOVA of the data with factor “groups” (Controls, Win-10, Win-20, Los-10, and Los-20) and one-way ANOVA of the data with factor “experience” of the groups (controls, Win-10, and Win-20) and (Controls, Los-10, and Los-20), followed by post hoc comparisons of the groups using the Bonferroni test or, for a number of repetitive behaviors, the LSD test. The data are reported as the mean ± SEM. A Chi-square analysis was used to identify significant differences in the percent of animals that exhibited attacks and aggressive grooming. The statistical significance was set at *P* ≤ 0.05. The experimental groups contained 10 mice in the control group and 11 mice in the Win-10, Win-20, Los-10, and Los-20 groups.

For the transcriptomic data, a Principal Components (PC) analysis was conducted using the XLStat software package (www.xlstat.com). It was based on a Pearson correlation metric calculated on the FPKM value profiles of 27 analyzed genes. Agglomerative Hierarchical Clustering (AHC) and Multidimensional Scaling (MDS) were performed on the same data with the XLStat software package. We also used a Pearson correlation as a similarity metric for the AHC/MDS analysis. The agglomeration method comprised an unweighted pair-group average. Multidimensional Scaling was performed by minimization of the Kruskal stress statistic, with a random initial configuration, 5 repetitions and stop conditions as follows: Convergence = 0.00001 / Iterations = 500.

The identification of alternatively spliced events in the RNA-Seq data was performed with rMATs software [48]. Refseq v. 10.0 was used as a template for mapping the reads and alternative event annotations. Only exon skipping events were considered.

## Results: Social interactions

### Avoidance behavior

One way ANOVA revealed a significant influence of the factor “groups” on the parameters of avoidance behavior: number (F (4, 48) = 23.64; *P* < 0.001) and total time (F (4, 48) = 33.07; *P* < 0.001) (Fig. 2). Based on the posthoc Bonferroni test as compared to the respective levels in the controls, the number of episodes and total time of avoidance behavior were increased in the Los-10 (P < 0.001 and *P* < 0.012, respectively) and in the Los-20 (for both *P* < 0.001). These parameters were significantly more in the Los-10 as compared with the Win-10 (P < 0.001 and *P* < 0.007, respectively) and in the Los-20 as compared with the Win-20 (for both P < 0.001).

**Figure 2.**
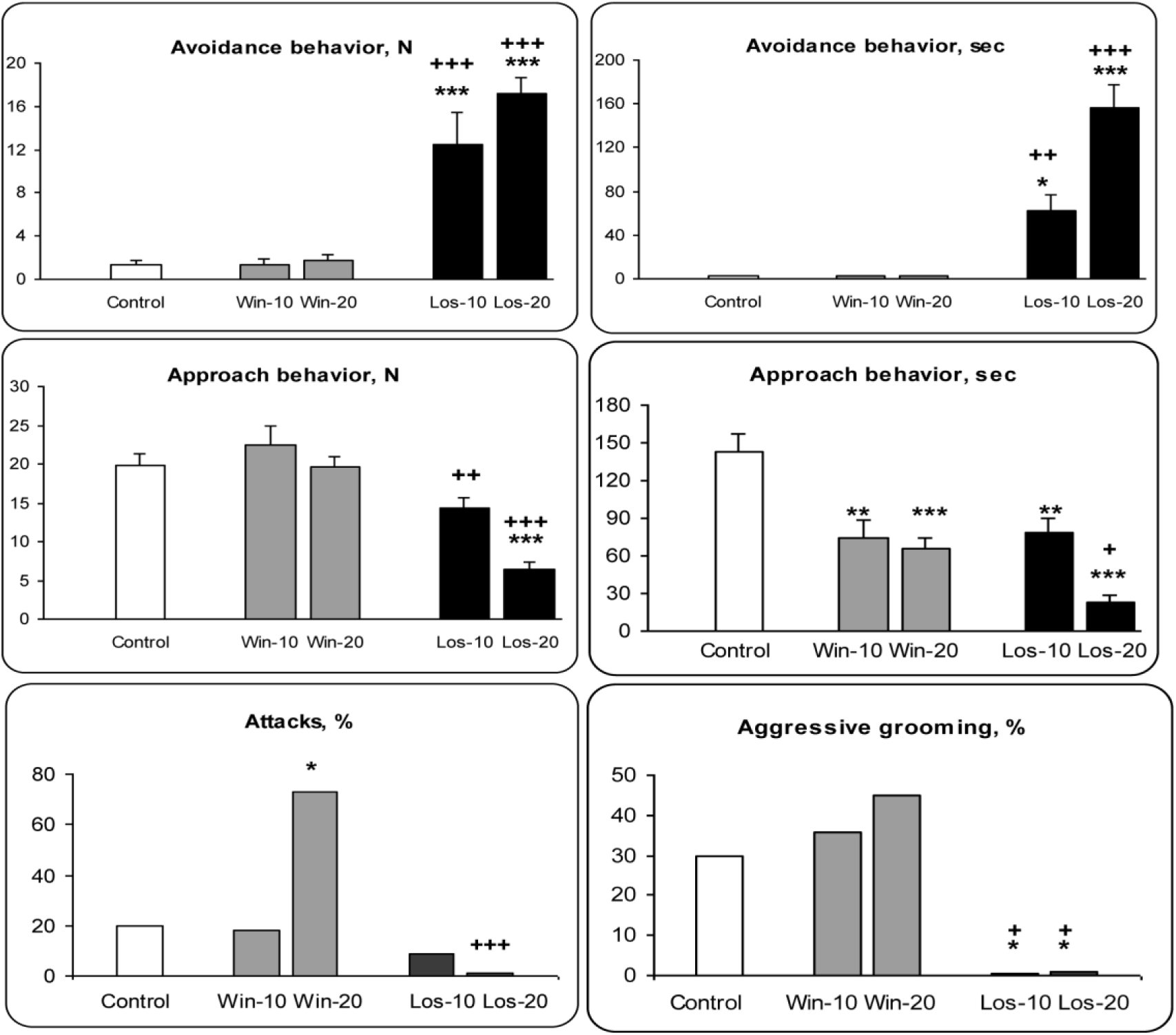
Social behaviors of the controls, winners and losers in the social interaction test. Los-10, Los-20 and Win-10, Win-20 - male mice with experiences of social defeats and wins, respectively, with 10 and 20 days of agonistic interactions. * - *P* < 0.05; ** - P < 0.01; *** - *P* < 0.001 *vs* controls; + - *P* < 0.05; ++ - *P* < 0.01; +++ - *P* < 0.001 *vs* winners of respective social experience. Significant differences between the groups of Win-10 *vs* Win-20 and Los-10 *vs*- Los-20 are described in the Results section.

One way ANOVA revealed a significant influence of the factor “experience” in the losers on the avoidance parameters: number (F (2, 28) = 15.51; *P* < 0.001) and total time (F (2, 28) = 24.24; *P* < 0.001). One way ANOVA did not revealed a significant influence of the factor “experience” in the winners (ns).

### Approach behavior

One way ANOVA revealed a significant influence of the factor “groups” on parameters of approach behavior: number (F (4, 48) = 16.05; *P* < 0.001) and total time (F (4, 47) = 13.21; *P* < 0.001) (Fig. 2). Based on the posthoc Bonferroni test, the number and total time of approach behavior were decreased in the Los-20 as compared with the respective levels in the controls (for both *P* < 0.001), and Win-20 (*P* < 0.001 and *P* < 0.014, respectively). Additionally the number of approaches was significantly less in the Los-10 as compared with the Win-10 (*P* < 0.007) and total time of approach behavior were decreased in the Los-10 (*P* < 0.004), Win-10 (*P* < 0.002), Win-20 (*P* < 0.001) in comparison with the controls.

One way ANOVA revealed a significant influence of the factor “experience” on the number of approaches (F (2, 28) = 29.52; *P* < 0.001) and total time (F (2, 27) = 30.92; *P* < 0.001) in the losers and on the total time (F (2, 27) = 10.01; *P* < 0.001) in the winners.

### Aggression and aggressive grooming

In the social interactions test the Win-20 demonstrated short aggression during 18,4 sec/per test on the average, and only one winner showed about 3 min of attacks and persecutions in total. Percent of the Win-20 demonstrating attacks was about 70% *vs* 20% in the controls (8/11 vs 2/10) as well as in the Los-10 (1/11) and Los-20 (0/11). There were no differences between proportion of the controls (3/10), Win-10 (4/11) and Win-20 (5/11), demonstrating aggressive grooming (Fig. 2). The losers never demonstrated aggressive grooming toward unfamiliar partner (0/11).

Chi-square analysis revealed differences in the number of animals demonstrating attacks between the controls and Win-20 (*P* = 0.016) and between the Los-20 and Win-20 (*P* = 0.001). Differences in the number of animals demonstrating aggressive grooming were revealed between the controls and the Los-10 and Los-20 (for both *P* < 0.05), between the Los-10 and Win-10 (*P* = 0.027) and between the Los-20 and Win-20 (*P* = 0.011).

### Rearing behavior

One way ANOVA revealed a significant influence of the factor “groups” on the number (F (4, 48) = 8.44; *P* < 0.001) and total time (F (4, 48) = 12.44; *P* < 0.001) of rearing behavior (Fig. 3). Based on the posthoc Bonferroni test as compared to the level in the controls, the number of rearing was decreased in the Los-10 (*P* < 0.001), Los-20 (*P* < 0.001) and Win-20 (*P* < 0.005) and total time of rearing display was decreased in the Win-10 (*P* < 0.008), Win-20, Los-10, Los-20, (for all *P* < 0.001).

One way ANOVA revealed a significant influence of the factor “experience” in the losers on the number (F (2, 28) = 21.40; *P* < 0.001) and total time (F (2, 28) = 26.19; *P* < 0.001) and in the winners on the number (F (2, 28) = 5.45; *P* < 0.01) and total time (F (2, 28) = 10.01; *P* < 0.001).

**Figure 3.**
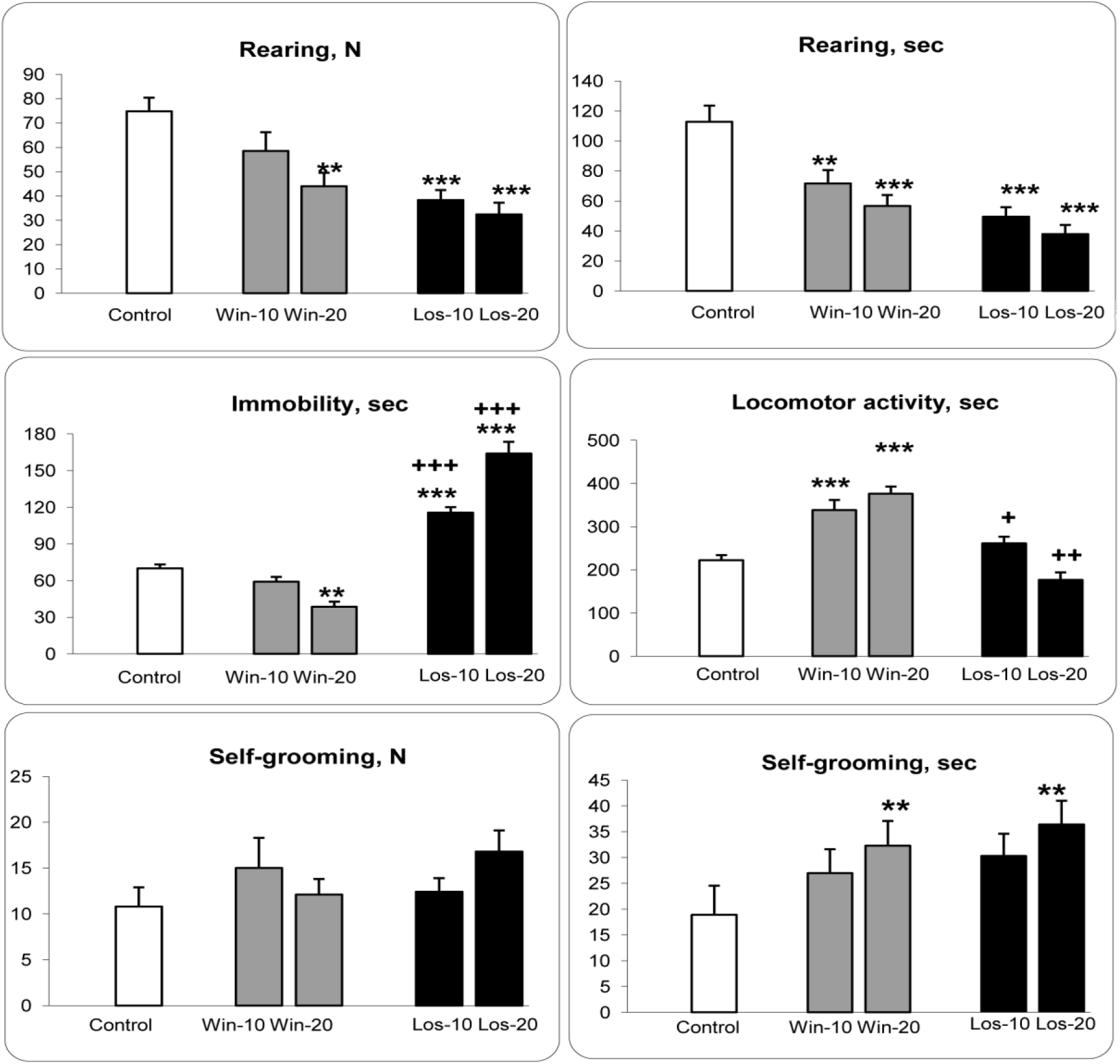
Individual behaviors of the controls, winners and losers in the social interaction test. Los-10, Los-20 and Win-10, Win-20 - male mice with experiences of social defeats and wins, respectively, with 10 and 20 days of agonistic interactions. * - *P* < 0.05; ** - *P* < 0.01; *** - *P* < 0.001 *vs* controls; + - *P* < 0.05; ++ - P< 0.01; +++ - *P* < 0.001 *vs* winners of respective social experience. Significant differences between the groups of Win-10 *vs* Win-20 and Los-10 *vs*- Los-20 are described in the Results section.

### Immobility

One way ANOVA revealed a significant influence of the factor “groups” on the total time of immobility (F (4, 48) = 78,71; *P* < 0.001) (Fig. 3). Based on the posthoc Bonferroni test as compared to the level in the controls, the total time was decreased in the Win-20 (*P* < 0.005) and was increased in the Los-10 and Los-20 (for both *P* < 0.001). Significant differences were shown in comparison of the Los-10 *vs* Los-20; Los-10 *vs* Win-10; Los-20 *vs* Win-20 (for all, *P* < 0.001). One way ANOVA revealed a significant influence of the factor “experience” in the losers (F (2, 28) = 47.02; *P* < 0.001) and in the winners (F (2, 28) = 16.42; *P* < 0.001).

### Locomotor activity

One way ANOVA revealed a significant influence of the factor “groups” on the total time of locomotion (F (4, 48) = 21.86; *P* < 0.001) (Fig. 3). Based on the posthoc Bonferroni test as compared to the controls, this parameter was increased in the Win-10 and Win-20 (for both *P* < 0.001). Significant differences were shown in comparison of the Los-10 *vs* Los-20 (P < 0.012); the Los-10 *vs* Win-10 (*P* < 0.029), in the Los-20 *vs* Win-20 (*P* < 0.001). One way ANOVA revealed a significant influence of the factor “experience” in the losers on (F (2, 28) = 7.81; *P* < 0.002) and winners (F (2, 28) = 17.59; *P* < 0.001).

### Self-grooming

One way ANOVA revealed a significant influence of the factor “groups” on the total time of self-grooming behavior (F (4, 47) = 2,77; *P* < 0.038) (Fig. 3). Based on the posthoc LSD test as compared to the level in the controls, total time was increased in the Win-20 (*P* < 0.008), Los-20 (*P* < 0.005) and Los-10 (*P* ≤ 0.05). One way ANOVA revealed a significant influence of the factor “experience” in the winners (F (2, 27) = 3.42; *P* < 0.047) and in the losers (F (2, 28) = 4.75; *P* < 0.017). In this comparison the Bonferroni test confirmed significant differences between the controls and Win-20 (*P* < 0.043) and the controls and Los-20 (*P* < 0.015). Figure 4 shows behaviors of the controls, Win-20 and Los-20 expressed as a percentage of total time for demonstration of locomotor activity, avoidance, approach behavior, rearing and immobility during social interactions test. On the neutral territory the controls demonstrated mainly exploration: locomotor activity and rearing time was about 60% of testing time. They displayed interest to an unfamiliar partner: the controls approached to unfamiliar male and spent many time sniffing it and never demonstrated avoidance behavior toward approaching partner. The winners also seldom demonstrated avoidance behavior. 70% of testing time they chaotically moved in the cage and did not pay any attention to the partner.

**Figure 4.**
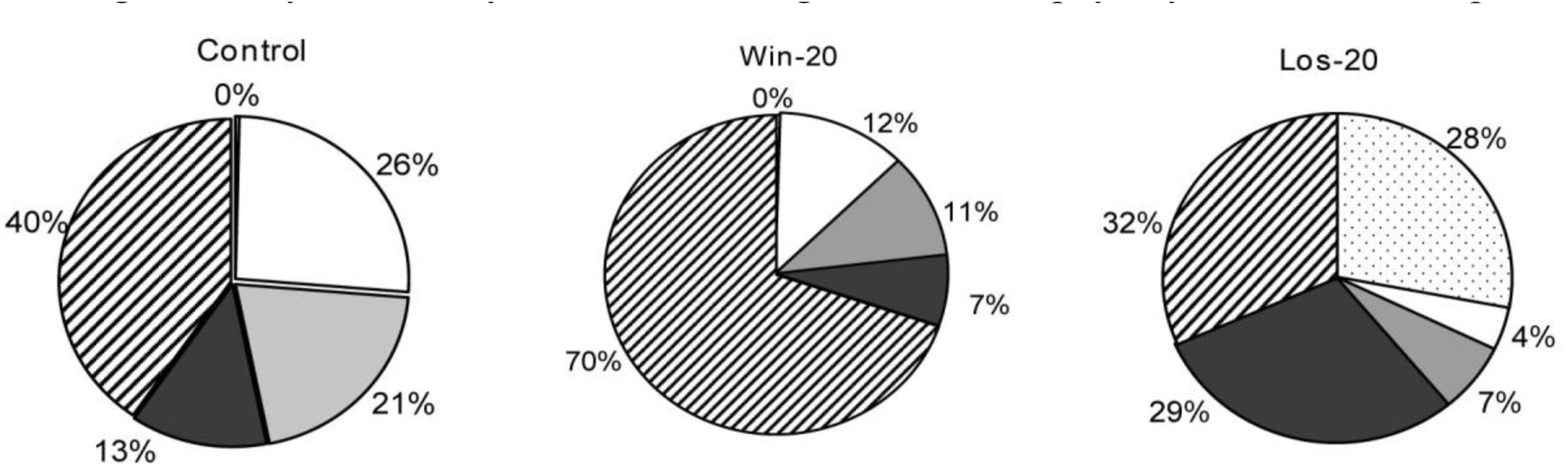
Behaviors of the controls, Win-20 and Los-20 expressed as a percentage of the total time for the demonstration of locomotor activity, avoidance, approach behavior, rearing and immobility during the social interaction test. 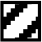 Locomotor activity; 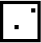 Avoidance; 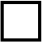 Approach behavior; 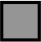 Rearing; 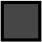 Immobility

Approaching the stranger, they attacked or displayed the aggressive grooming behavior. The losers actively avoided an amicable stranger or demonstrated immobility behavior about 58% of testing time. They also seated in the corner or near a wall of cage, watched the partner and never displayed aggressive behavior toward him. They rarely themselves approached to an unfamiliar male and sniffed it as the control mice did.

### Partition test

One way ANOVA revealed a significant influence of the factor “groups” on the number of approaches (F (4, 47) = 7.47; *P* < 0.001) and total time (F (4, 47) = 40.18; *P* < 0.001) parameters (Fig. 5). Based on the posthoc Bonferroni test as compared to the level in the controls, number of approaches was decreased in the Los-10 (*P* < 0.001) and Los-20 (*P* < 0.003); significant differences were shown in comparison of the Los-10 *vs* Win-10 (*P* < 0.034) and the Los-20 *vs* Win-20 (*P* ≤ 0.051). Total time spent near the partition differs significantly in comparison of the control *vs* Los-10 and Los-20 (for both *P* < 0.001) and *vs* the Win-10 (*P* < 0.001); the Los-10 *vs* Win-10 (P < 0.001) and in Los-20 *vs* Win-20 (*P* < 0.001). One way ANOVA revealed a significant influence of the factor “experience” in the losers on number of approaches (F (2, 28) = 9.84; *P* < 0.001) and total time (F (2, 28) = 34.30; *P* < 0.001) and in the winners on total time (F (2, 27) = 7.79; *P* < 0.002).

**Figure 5.**
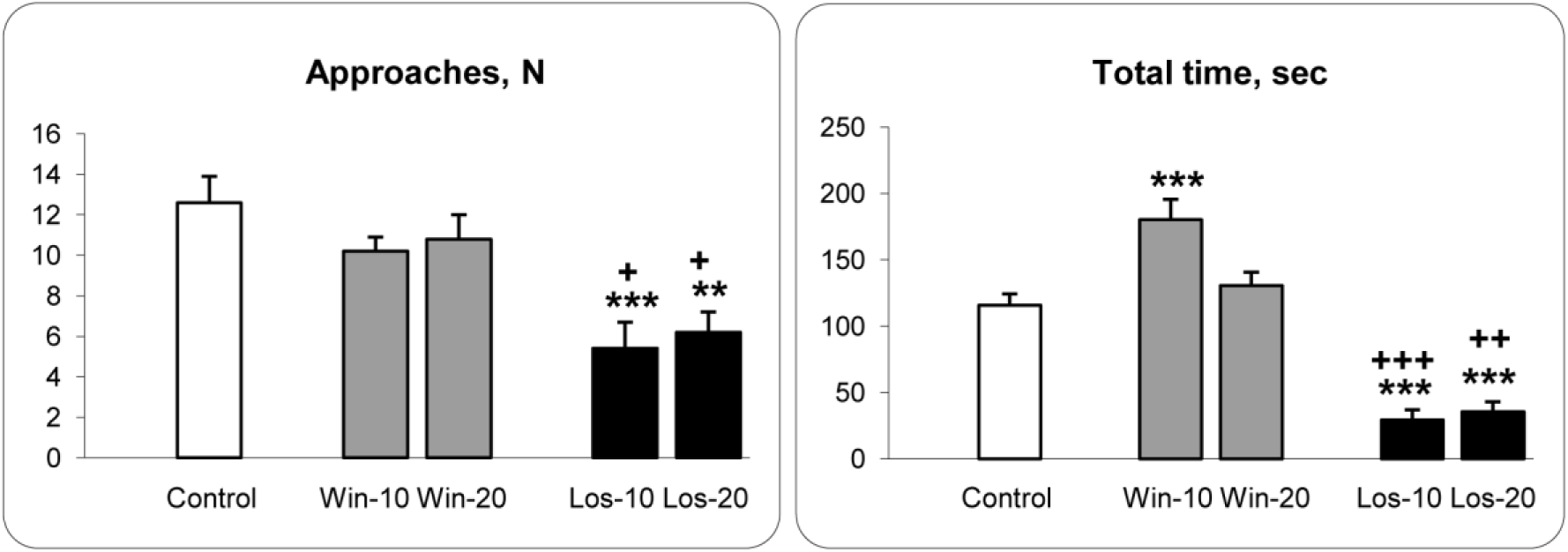
Behaviors of the controls, winners and losers in the partition test. Los-10, Los-20 and Win-10, Win-20 - male mice with experiences of social defeats and wins, respectively, with 10 and 20 days of agonistic interactions. ** - *P* < 0.01; *** - *P* < 0.001 *vs* controls; + - *P*< 0.05; +++ - *P* < 0.001 *vs* winners of respective social experience.

## Results: RNA-Seq study

Analyzing of whole transcriptome data of the midbrain raphe nuclei, ventral tegmental area (VTA), striatum, hippocampus, and the hypothalamus, we found in the winners and losers common and different changes in the expression of autism-related genes of interest. Data of the RNA-Seq in FPKM values for differentially expressed 27 genes in brain regions are presented in Supplementary Table 1, 2 (at the end).

In the hypothalamus **(Fig. 6a,b)** autism-related genes decreased their expression both in the winners and losers in comparison with the controls as for genes *Maoa* (*P* ≤ 0.020 and *P* < 0.0001; q < 0.05, respectively), *Gabrb3* (*P* ≤ 0.038 and *P* < 0.05, respectively), *Fmr1* (*P* ≤ 0.001; q < 0.05 and *P* < 0.003; q < 0.05, respectively), *Ube3a* (*P* < 0.038 and *P* < 0.003; q < 0.05, respectively), *Pten* (*P* ≤ 0.0023; q < 0.05 and *P* < 0.0002; q < 0.05, respectively), *Mecp2* (*P* ≤ 0.039 and *P* < 0.05, respectively), *Ptchd1* (*P* < 0.0008; q < 0.05 and *P* < 0.0001; q < 0.05, respectively), and *Nrxn1* (*P* ≤ 0.043 and *P* < 0.013, respectively). In both social groups expression of the *Shank3* gene was increased (*P* ≤ 0.003 and *P* < 0.0001; q < 0.05, respectively) (Fig. 6a,b).

**Figure 6.**
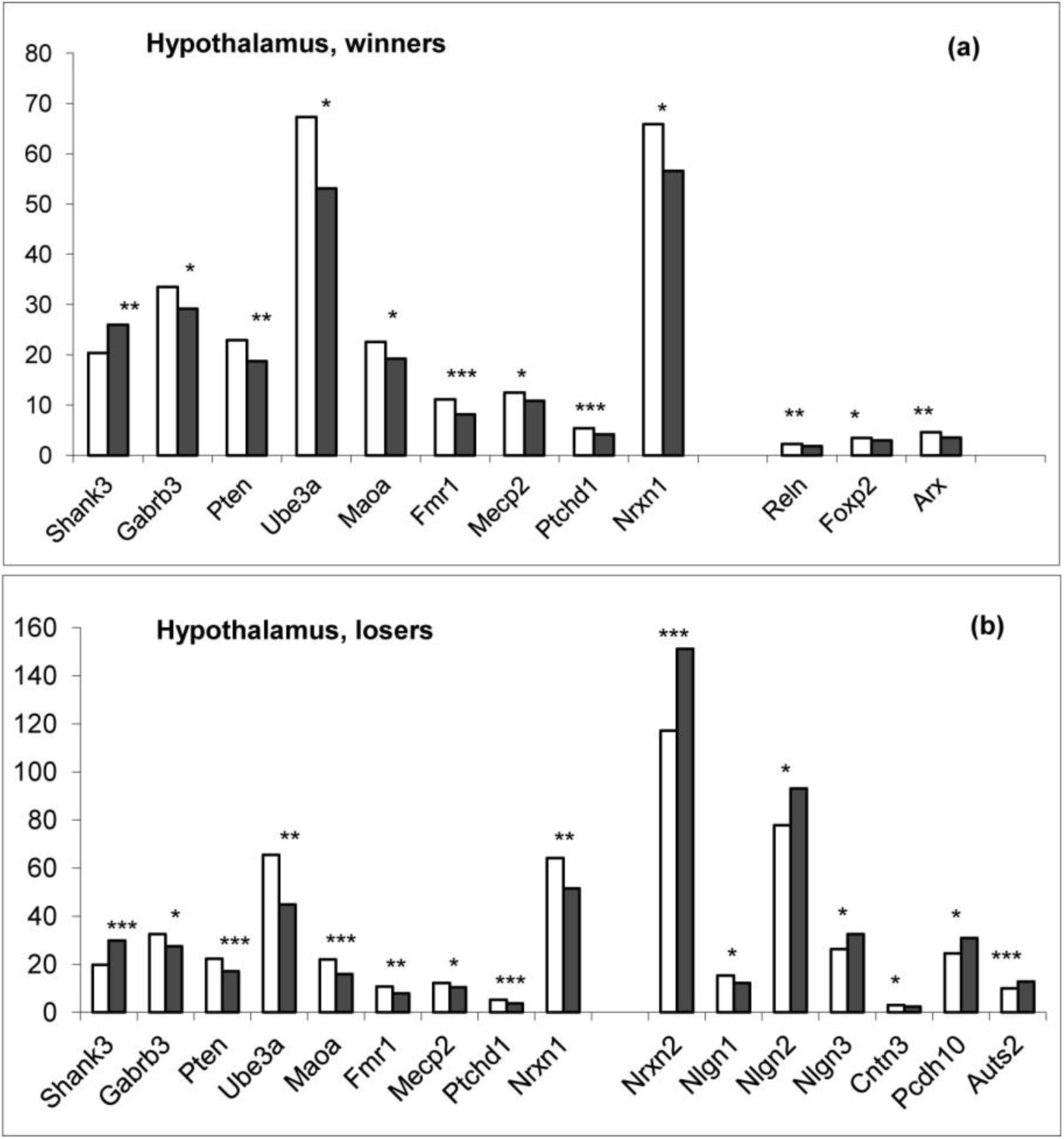
Differentially expressed autism-related genes in the hypothalamus of the winners (a) and losers (b) under agonistic interactions. The Cufflinks program was used to estimate the gene expression levels in FPKM units. White columns – the control; black columns – winners or losers. * - *P* < 0.05; ** - *P* < 0.01; *** - *P* < 0.001 *vs* controls.

In comparison with the controls, in the winners expressions were decreased in the *Reln* (*P* < 0.009), *Foxp2* (*P* < 0.037), and *Arx* (*P* < 0.006) genes. In the losers, expression of the *Nrxn2* (*P* < 0.0012; q < 0.05), *Nlgn2* (*P* ≤ 0.022), *Nlgn3* (*P* ≤ 0.014), *Auts2* (*P* < 0.001; q < 0.05), and *Pcdh10* (*P* < 0.020) genes were increased and of the *Nlgn1* (*P* < 0.020) and *Cntn3* (*P* ≤ 0.017) genes were decreased. In common, 12 autistic genes in the winners and 16 genes in the losers changed their expression under agonistic interactions.

In the midbrain raphe nuclei expression of the *Tph2* (for both groups *P* < 0.0001; q < 0.05), *Slc6a4* (for both groups *P* < 0.0001; q < 0.05), *Foxp2* (for both groups *P* < 0.0001; q < 0.05), *Reln* (for both groups *P* < 0.0001; q < 0.05), and *Oxtr* (*P* < 0.003 for the winners and *P* < 0.010 for the losers) genes is decreased in both social groups (Fig. 7). Additionally in the winners expression of the *Ube3a* (*P* < 0.035), *Fmr1* (*P* < 0.042), *Nrxn1* (*P* < 0.048), *Cadps2* (*P* < 0.034), and *Cntn3* (*P* < 0.024) genes were increased. There were no specific changes in this brain region in the losers. In common, 10 genes in the winners and 5 genes in the losers changed their expression in the midbrain raphe nuclei under agonistic interactions.

**Figure 7.**
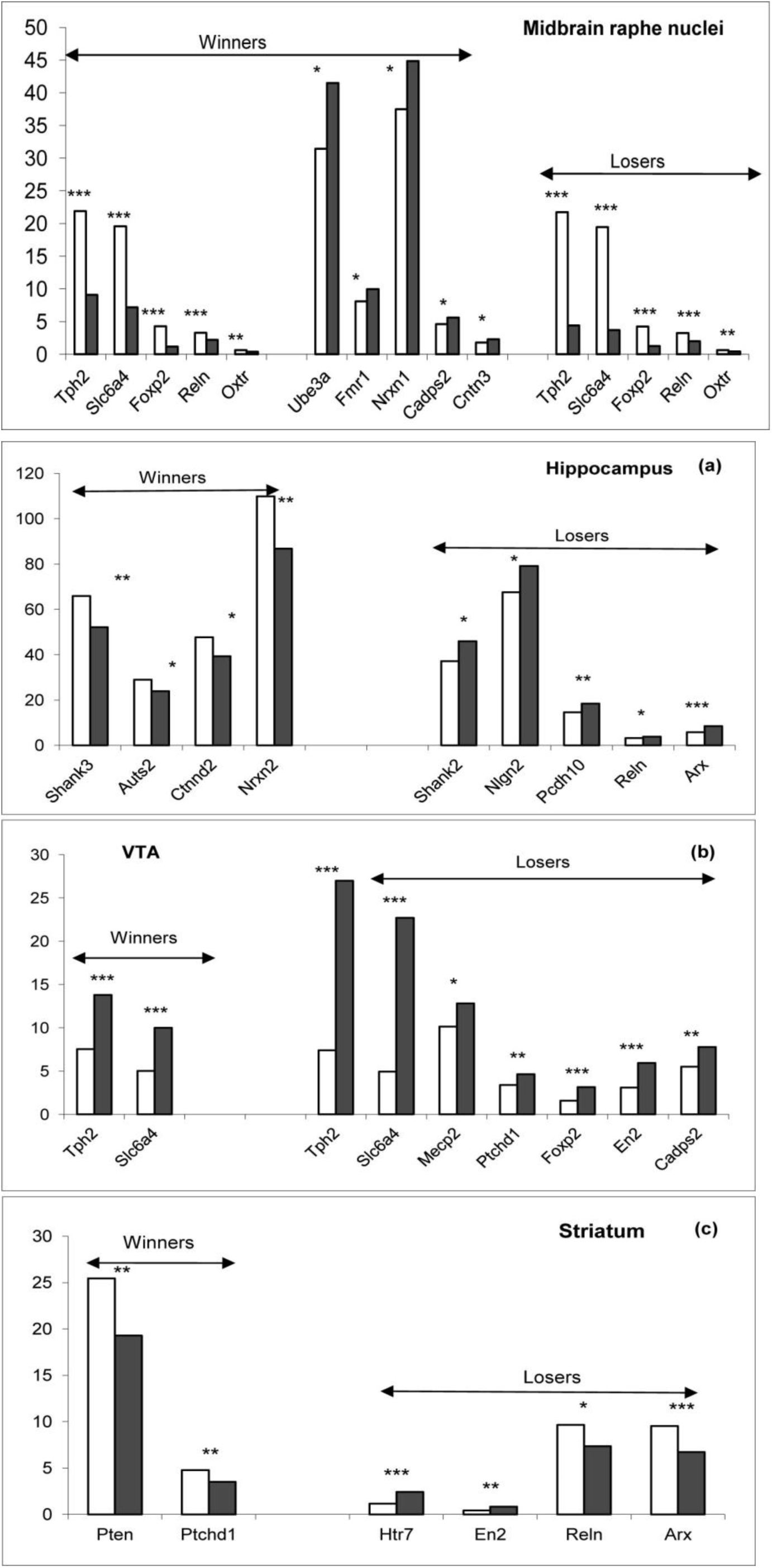
Differentially expressed genes in the midbrain raphe nuclei of the winners and losers. The Cufflinks program was used to estimate the gene expression levels in FPKM units. White columns – the control; black columns – winners or losers. * - *P* < 0.05; ** - *P* < 0.01; *** - *P* < 0.001 *vs* controls.

In the hippocampus there are no common for both social groups genes that changed their expression. However the reduced expression of the *Shank3* (*P* < 0.010), *Auts2* (*P* < 0.024), *Nrxn2* (*P* ≤ 0.011) and *Ctnnd2* (*P* < 0.021) genes was revealed in the winners. The *Shank2* (*P* < 0.040), *Nlgn2* (*P* < 0.047), *Pcdh10* (*P* ≤ 0.011), *Reln* (*P* < 0.026) and *Arx* (*P* < 0.0002; q < 0.05) genes increased their expression in the losers (Fig. 8a). In common, 9 autistic genes (4 genes in the winners and 5 genes in the losers) changed their expression in the hippocampus under agonistic interactions.

**Figure 8.**
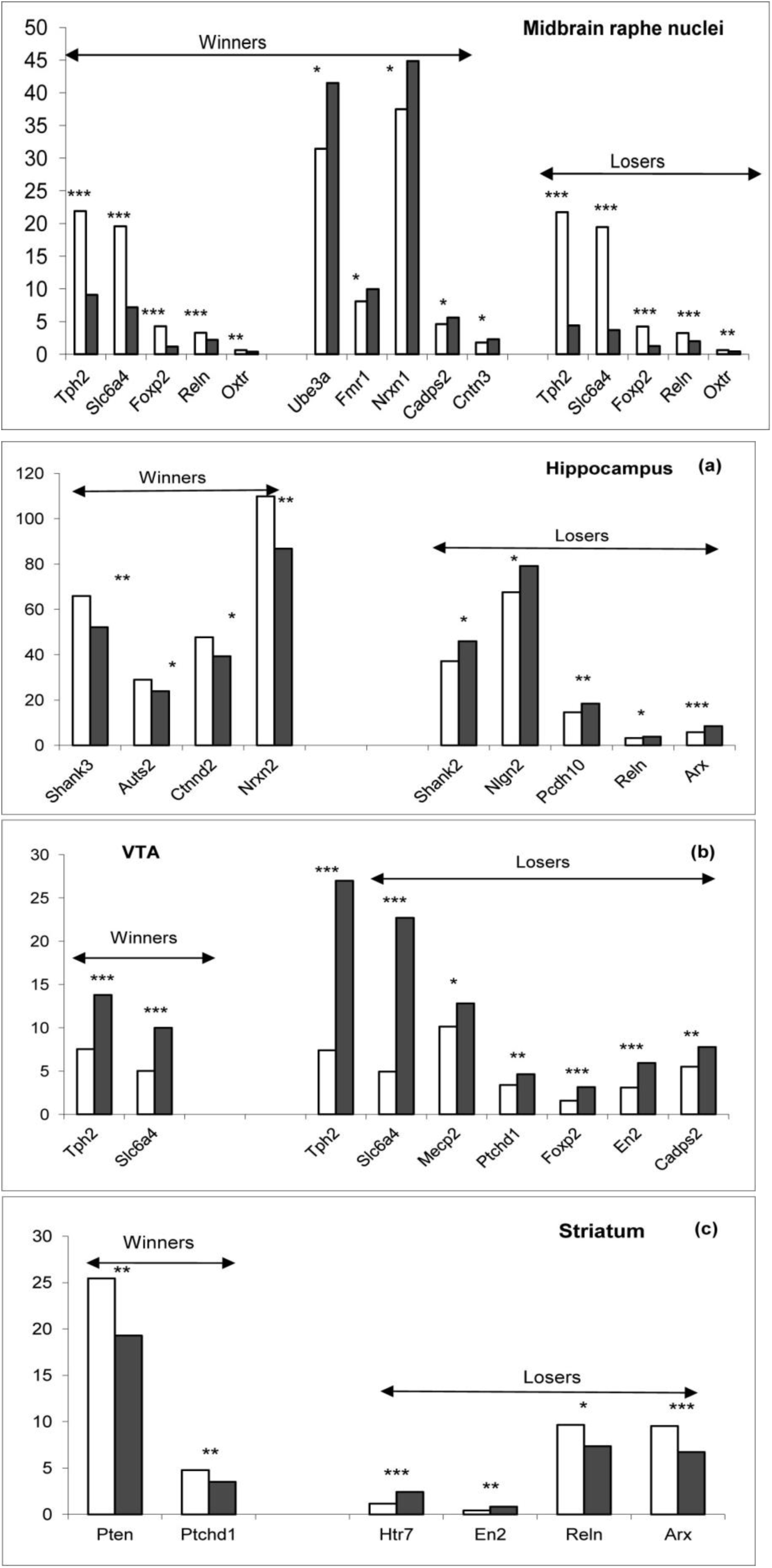
Differentially expressed genes in the hippocampus (a), VTA (b) and striatum (c) of the winners and losers under agonistic interactions. The Cufflinks program was used to estimate the gene expression levels in FPKM units. White columns – the control; black columns – winners or losers. * - *P* < 0.05; ** - *P* < 0.01; *** - *P* < 0.001 *vs* controls.

In the VTA increased expression of the *Tph2* (*P* < 0.0001; q < 0.05) and *Slc6a4* (*P* < 0.0001; q < 0.05) genes were common for both social groups. The *Mecp2* (*P* < 0.042), *Ptchd1* (*P* < 0.012), *Foxp2* (*P* < 0.0001; q < 0.05), *En2* (*P* < 0.0001; q < 0.05) and *Cadps2* (*P* < 0.005) genes were upregulated only in the losers (Fig. 8b).

In the striatum the *Pten* (*P* < 0.010) and *Ptchd1* (*P* < 0.010) genes in the winners and the *Reln* (*P* ≤ 0.033) and *Arx* (*P* ≤ 0.020) genes were downregulated in the losers under repeated agonistic interactions (Fig. 8c). In the losers, expression of the *Htr7* (*P* < 0.0001; q < 0.05) and *En2* (*P* < 0.011) genes were upregulated.

Expression of the *Bdnf* gene was increased in the striatum (*P* < 0.0001; q < 0.05) of the losers and was decreased in the midbrain raphe nuclei in the winners and losers (*P* ≤ 0.0015, *P* ≤ 0.0023, respectively).

### Concordant and discordant changes in gene expression in the winners and losers

In the hypothalamus of the winners and losers, the expressions of the *Maoa, Mecp2, Ube3a, Gabrb, Cntn3, Pten, Nrxn1, Fmr1*, and *Ptchd1* genes were downregulated and the *Shank3* gene was upregulated. In the midbrain raphe nuclei of both social groups, the *Tph2, Slc6a4, Reln, Foxp2*, and *Oxtr* genes were downregulated. In the VTA, the *Tph2* and *Slc6a4* genes were upregulated.

Specific changes for the winners included the downregulation of the *Reln, Foxp2* and *Arx* genes in the hypothalamus, the *Shank3, Auts2, Ctnnd2*, and *Nrxn2* genes in the hippocampus, and the *Pten* and *Ptchd1* genes in the striatum. In contrast, the *Ube3a, Fmr1, Nrxn1, Cadps2*, and *Cntn3* genes were upregulated in the midbrain raphe nuclei of the winners.

Specific changes for the losers included the upregulation of the *Shank2, Nlgn2, Pcdh10, Reln*, and *Arx* genes in the hippocampus; the *Mecp2, Ptchd1, Foxp2, En2*, and *Cadps2* genes in the VTA; the *Nrxn2, Nlgn2, Nlgn3, Pcdh10*, and *Auts2* genes in the hypothalamus; and the *En2* and *Htr7* genes in the striatum. An opposite downregulation was identified for the *Ube3a* and *Nlgn1* genes in the hypothalamus and the *Reln* and *Arx* genes in the striatum of the losers.

### Principal components analysis of autistm-related gene expression in the RNA-Seq data

To assess the degree of brain region-specific expression of genes of interest, we performed a Principal components analysis based on the co-variation of 27 genes using the expression profiles of 45 samples, which comprised RNA-Seq FPKM data for 5 brain regions of 9 mice. Ovals correspond to brain clusters. We identified compact clustering of the hypothalamus (HPT), striatum (STR) and hippocampus (HPC) samples based on gene expression profiles (Fig. 9, encircled), whereas the midbrain raphe nuclei (MRN) and ventral tegmental area (VTA) were not distinct.

**Figure 9.**
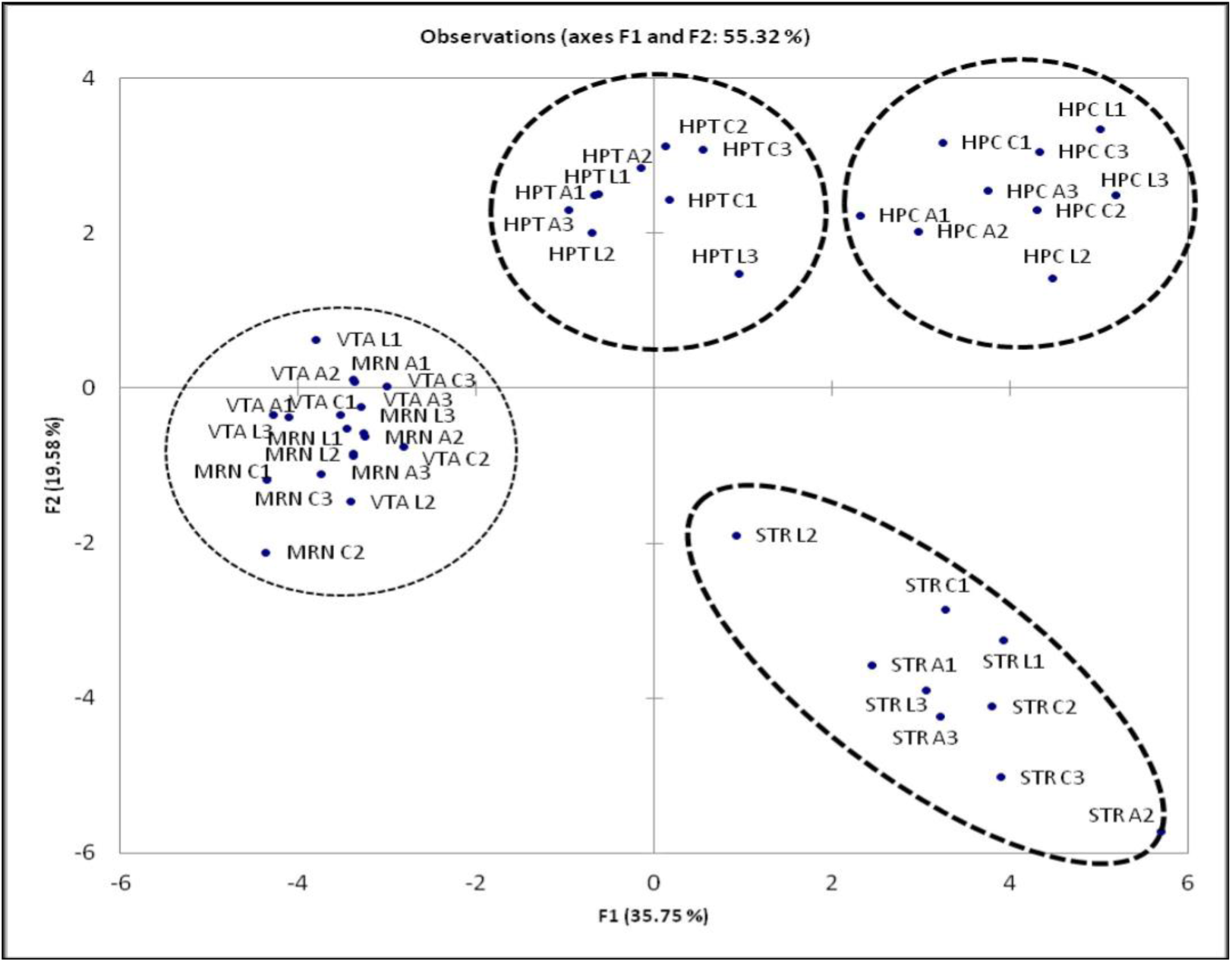
Principal Component (PC) analysis plot based on co variation of 27 autistic genes using the expression profiles of 45 samples, which comprised RNA-Seq FPKM data for 5 brain regions of 9 mice. Ovals correspond to brain regions. Abbreviations: HPT – Hypothalamus; HPC – Hippocampus; MRN – Midbrain Raphe Nuclei; VTA – Ventral Tegmental Area; STR – Striatum; C (C1, C2, C3) – control; A (A1, A2, A3) – aggressive mice; L (L1, L2, L3) – defeated mice, losers. Distinct clustering of three brain regions occurred, whereas the MRN and VTA brain compartments were not distinct.

### Agglomerative hierarchical clustering of autism-related coding transcripts and identification of outliers

We applied agglomerative hierarchical clustering (AHC) to the initial sample of 27 genes across 45 samples, which comprised RNA-Seq FPKM data for 5 brain regions of 9 mice based on their expression profiles. Fig. 10 presents the dendrogram of gene clustering. The similarity ordinate corresponds to the Pearson correlation coefficient (df=29) (Supplementary Table 3).

**Figure 10.**
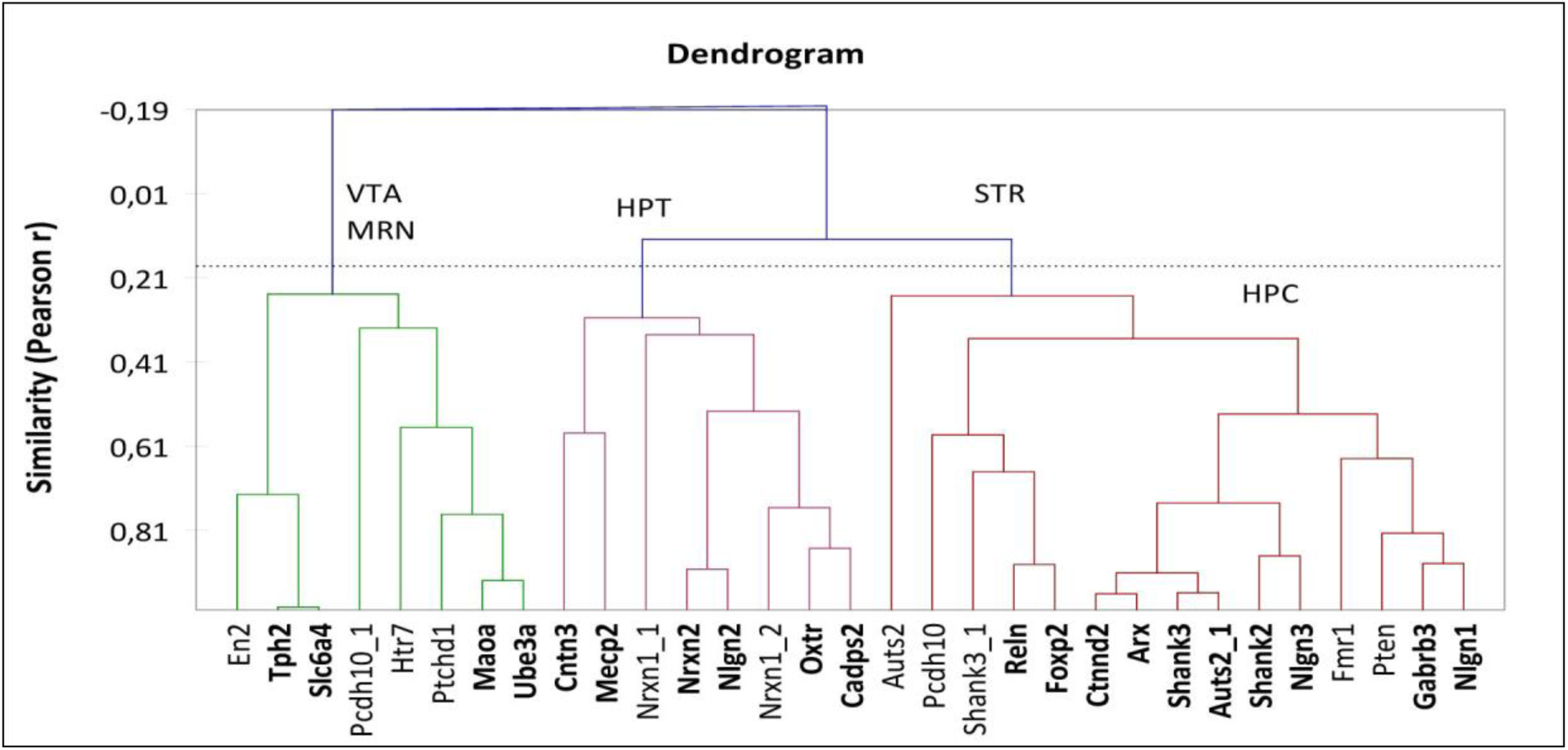
Agglomerative Hierarchical Clustering (AHC) tree for 27 autistic genes across 45 samples, which comprised RNA-Seq FPKM data for 5 brain regions of 9 mice. The genes elucidated as differentially expressed compared with the control, aggressive mice and defeated mice are marked in bold type script. The similarity ordinate corresponds to the Pearson correlation coefficient (df=29). Several clusters specific for the brain regions were elucidated. Abbreviations: HPT – Hypothalamus; HPC – Hippocampus; MRN – Midbrain Raphe Nuclei. Based on the expression, we determined the clusters as HPC-altered (typically down); STR-altered; HPT-altered; and MRN- and VTA- altered cluster.

An AHC analysis elucidates closely correlated genes, which thus strengthens the confidence of concordant differential expressions of a gene pair based on their highly correlated expression profiles across 45 samples (Fig. 10). Thus, we reaffirm the significant differential expression given correlated pairs of genes and independent tests using the Cufflinks program. For example, the *Tph2* and *Slc6a4* genes both exhibit downregulation in the midbrain raphe nuclei and upregulation in the VTA of the winners and losers while maintaining the co-expression correlation value across 45 samples, r=0.99; df=29 (Supplementary Table 3); the *Maoa* and *Ube3a* genes (r=0.93) and the *Cntn3* and *Mecp2* genes (r=.058) exhibit downregulation in the hypothalamus of the winners and losers; both the *Nrxn2* and *Nlgn2* genes (r=0.90) exhibit upregulation in the hypothalamus in the losers [which correspond to inhibitory synapses; 49]; the *Oxtr* gene exhibits downregulation and the *Cadps2* gene exhibits upregulation in the midbrain raphe nuclei of the winners (r=0.85); the *Reln* and *Foxp2* genes (r=0.89) exhibit downregulation in the midbrain raphe nuclei of the winners and losers; the *Ctnnd2* and *Arx* genes (r=0.96) exhibit downregulation in the hypothalamus and hippocampus in the winners; the *Shank3* and *Auts2* (r=0.62) genes exhibit downregulation in the hippocampus of the winners and upregulation in the hypothalamus of the losers; the *Shank2* gene exhibits upregulation in the hippocampus and the *Nlgn3* gene (r=0.87) exhibits upregulation in the hypothalamus of the losers; and the *Gabrb3* and *Nlgn1* (r=0.89) genes exhibit downregulation in the hypothalamus in the losers.

### Multidimensional Scaling (MDS) analysis of 27 neurospecific target genes and comparison with the Strings gene association network

We reconstructed the MDS plot based on the initial sample of 27 genes across 45 samples, which comprised RNA-Seq FPKM data for 5 brain regions of 9 mice based on their expression profiles. Figure 11 presents the gene relations based on the Pearson pairwise correlation coefficient values. We identified clustering of synapse versus serotonergic genes as indicated in the Strings gene network (Fig. 1). Intriguingly, in the comparison of the Strings network and MDS plot, we identified discrepancies as a result of the high negative correlation values, which thus misplaced the genes in the MDS plot while maintaining close relations. For example, the *Gabrb3* gene exhibits a distinct negative correlation with the serotonergic genes cluster as follows: *Maoa* (r=-0.25), *Tph2* (r=-0.53), *Slc6a4* (r=-0.5) and *Htr7* (r=-0.66) based on our RNA-Seq data (Supplementary Table 3); thus, it is positioned distantly from these four genes in Figure 11 in contrast with Figure 1. This finding implies that the Strings network (Fig. 1), while presenting the actual bibliographic association of genes, does not elaborate it in terms of the sign; however, it is based only on the joint occurrence in the same publications.

**Figure 11.**
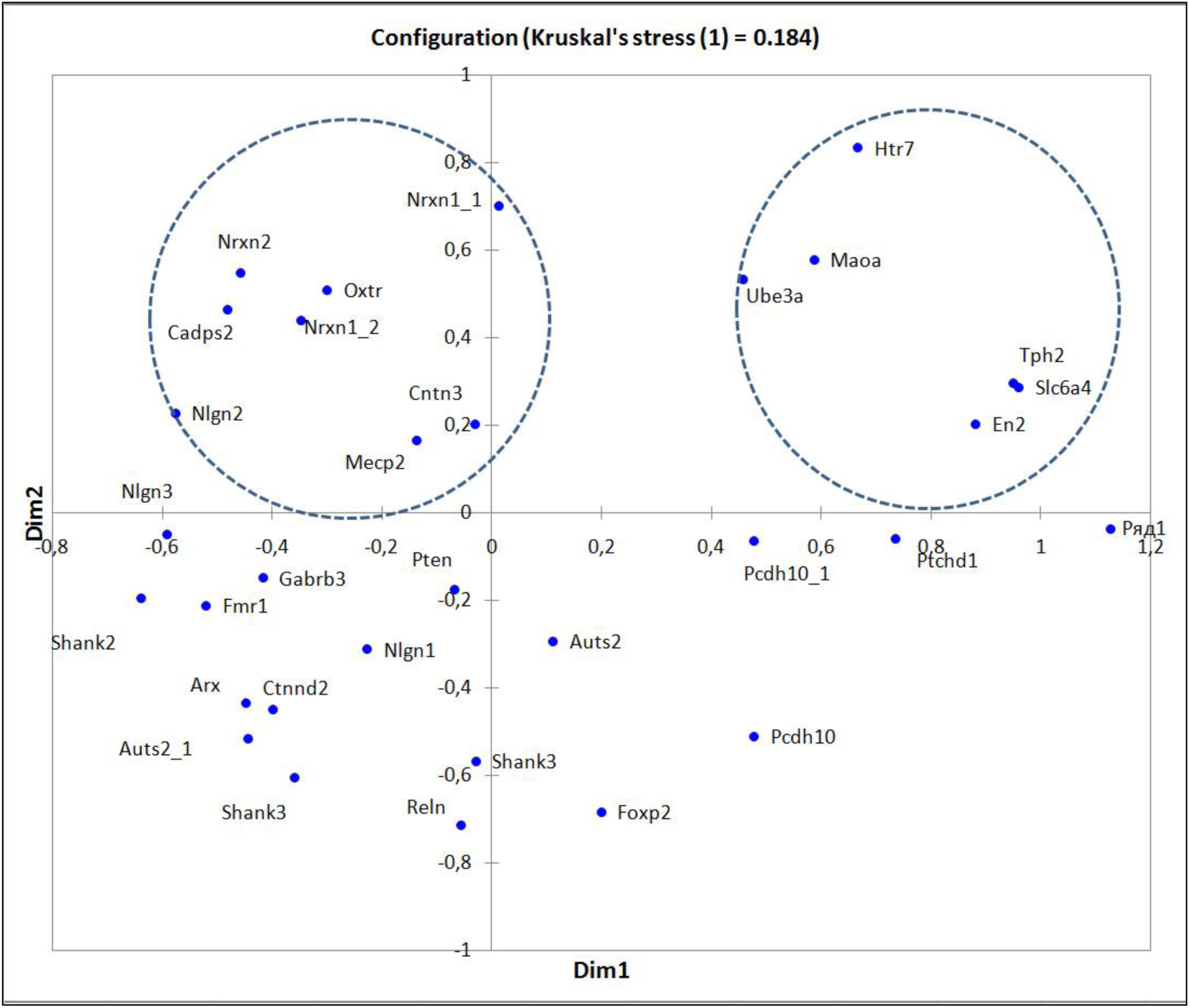
Multidimensional Scaling (MDS) plot of RNA-Seq co-variation data. Left circle corresponds to synaptic genes, and right circle corresponds to the serotonergic system.

### Alternative splicing

Seventeen of 27 autism related genes maintain alternatively spliced isoforms (Supplementary Table 1; Fig. 1). The major splice variability is provided by synapse specialized genes as previously demonstrated in a range of publications [49]. The largest number of isoforms observed in our data maintains the *Nrxn* (10 isoforms) cell adhesion molecule as follows: two isoforms identified in 2 splice variants were observed. Notably, the isoforms are distinctly expressed in particular brain regions as deduced from the PC analysis for the *Nrxn* and *Nlgn* gene families.

## Discussion

### Abnormal social behaviors associated with autistic spectrum symptoms induced by chronic agonistic interactions

This study demonstrates that the behaviors of male mice with alternative types of social behavior, the winners and losers, significantly differed in the social interactions test on neutral territory compared with the controls animals. Mice with repeated experiences of social defeats actively avoided approaching an amicable stranger or demonstrated freezing behavior, which may be considered passive avoidance of social contact with other partner on neutral territory. Most of the time, the defeaters sat in the corner or near a wall of the cage, watched for the partner and never demonstrated aggressive behavior or aggressive grooming toward him. They rarely approached an unfamiliar male and sniffed it compared with the control mice.

The total time of self-grooming behavior, which we initially considered a replacement activity, was increased in the losers. Similar to previous studies [29, 33, 35], the losers also exhibited low interest in the partner in the home cage as follows: the number of approaches and total time spent near the partition as a reaction to the partner in the neighboring compartment were significantly decreased. A low response to a stranger may be interpreted as low communication and an avoidance of social interactions.

Winners with repeated experience of aggression very seldom demonstrated avoidance behavior toward a forthcoming unfamiliar partner. They approached him; however, the total time of following for the partner was decreased because the winners, as a rule, attacked or exhibited aggressive grooming behavior. For most of the 10 min test, approximately 5-6 min, the winners chaotically moved in the cage and did not pay attention to the partner as follows: the total time of locomotion activity was significantly increased compared with the losers. It may be considered a development of hyperactivity and attention deficit behavior. The total time of self-grooming episodes was increased in the winners. In the partition test, the reaction to the partner in the neighboring compartment was increased in the Win-10. This behavior has previously been connected with aggressive motivation because behavioral activity near the partition measured prior to an agonistic interaction has been positively correlated with the total time of attacks, which winners demonstrated after the partition was removed [29, 34].

Decreased exploratory activity was identified both in the winners and losers as follows: the number and total time of rearing events were less compared with this parameter in the controls. In the winners, the exploration time decreased as a result of increased locomotion. In the losers, the exploration time decreased as a result of the increased immobility time. The different behavioral motivations may also be considered for the decreased approaching behavior time, which may reflect a deficit in communication or abnormal social behavior. As previously discusses, the repetitive self-grooming behaviors increased in both the Win-20 and Los-20 compared with the controls. An ANOVA indicated the influence of social experience duration (days) on the expression of most behavioral parameters in both social groups. As a rule, the severity of the behavioral changes was greater in the male mice with 20-day agonistic interactions.

Careful consideration of these results enabled us to note the intriguing similarities in the social behaviors and psychoemotional state of the winners and losers with several symptoms that other authors have considered autistic-like behaviors in animal models of autism [25, 27, 28, 50]. Impairments in social recognitions were common in both animal social groups. Experienced winners lose the ability to differentiate sex, age and social status of conspecifics and, as a rule, demonstrate irritability and aggression toward a partner even in low provoking situations [34]. They distantly differentiated males from receptive females substantially worse than the controls and equally attacked the other males that demonstrated full submission or active defense [31, 51], as well as young males [52]. We suggest that this behavior may be considered low empathic and aberrant social behavior. The losers exhibit avoidance behavior and immobility toward any partners. They differentiated familiar and unfamiliar conspecifics in the partition test significantly worse than the controls and exhibited full indifference, helplessness and absence of interest to novelty [30, 33, 53]. In addition to self-grooming behavior, the winners demonstrated different forms of other stereotypic repetitive behaviors, such as jumps, rotations, back circles, head shakes, and rigid tail, in other behavioral tests [34, 54]; however, the expression and types of stereotypes may be different in the winners of different strains. An increased number of crossed squares (hyperactivity) in the open field test was identified in the winners.

Enhanced anxiety, which did not disappear two weeks after stopping agonistic interactions [55, 56], was demonstrated in both social groups. Notably, according to different reports, anxiety disorders are present in 11% - 84% of children with autistic symptoms [57-60]. Moreover, similar to ASD patients [61], both the losers and winners exhibited general reward dysfunction. For example, they decreased sucrose solution consumption in a free two-bottle choice paradigm (34, 62) as a result of disturbances in motivated rewarding behaviors, as well as the prevalence of anxiety, fear and depressiveness in the losers and anxiety and aggressive motivation in the winners. Thus, all main symptoms and associated behaviors indicate the development of autistic symptoms during long-term agonistic interactions in male mice (Table 1). Thus, the behavioral data demonstrated similarities of autistic spectrum symptoms and changes in social and associated behaviors in animals with repeated experiences of agonistic interactions.

**Table 1.** ASDs symptoms in humans, their animal analogs in gene-modified animals and in the aggressive and defeated mice

|  |  |  |  |
| --- | --- | --- | --- |
| Symptoms in humans<br>DSM-4, DSM-5 | Symptoms in animal<br>models* | Winners** | Losers*** |
| **Main symptoms** |  |  |  |
| Low communication | Reduced or low sociability | Reduced or low sociability | Reduced or low sociability<br>Avoidance of social contact |
| Impaired socialization | Impaired social recognition<br>Cognitive deficit | Impaired social recognition<br>Cognitive deficit | Impaired social recognition<br>Cognitive deficit |
| Restricted and/or repetitive behavior, seizures, stereotypies, dyskinesia | Increased repeated motor activities (spontaneous activity, exploration, circling, digging, jumping)<br>Increased self-involved behaviors (self-grooming, scratching, circling behavior) | Increased repetitive self-grooming, jumping, circling, jerk, digging, movement back, rotations, head shakes *etc.* | Increased repetitive self-grooming |
| **Associated symptoms** |  |  |  |
| Attention/deficit |  | Attention/deficit | Social indifference |
| Hyperactivity |  | Hyperactivity, hyperexcitation | Hypoactivity (freezing, immobility, helplessness, depressiveness) |
| Hyperresponsiveness<br>Hyporesponsiveness |  | Hyperresponsiveness to sensory stimuli | Hyporesponsiveness to sensory stimuli |
| Anxiety, depression | Increased anxiety-like behavior | Increased anxiety | Increased anxiety, depression |
| Aggression, irritability |  | Aggression, irritability | Social avoidance |
| Exploration deficit | Exploration deficit | Exploration deficit | Exploration deficit |
| Reward dysfunction |  | Reward dysfunction | Reward dysfunction |
| * - [25-28]; ** - [3, 54, 55]; these data; *** - [32, 30, 34, 62]; these data. |  |  |  |

The question arises as to whether the changes in communication and sociability, as well as in associated social behaviors and the psychoemotional state in the winners and losers may be regarded as ASD–like symptoms. According to the DSM-V [2], the diagnostic criteria of autism include the following triad of symptoms: impaired social interactions, low communication and repetitive behaviors. These basic symptoms may be accompanied by restricted interests and altered activity, atypical motivated behavior [61], anxiety disorders [58–60], enhanced aggressiveness and irritability, and attention/deficit hyperactivity [11, 63]. Moreover, the social behaviors of chronically defeated and aggressive mice are essentially indicative of all symptoms of ASDs (Table 1), which have been used in models of autism, such as in animals of specific strains, transgenic or knockout mice [25, 27, 28, 50].

However, this conclusion did not meet understanding in the majority of researchers working in the field of autism modeling in animals. There is a common representation that autism and ASDs are developmental disorders, which are particularly present in childhood and are genetically determined. To confirm or reject our observation, we conducted a study of the expression of genes that are involved in the mechanisms of autism. We identified numerous changes in the expression of recognized autism genes in animals with abnormal social behaviors. We demonstrate changes in gene expression in different brain regions of male mice with repeated experiences of aggression and social defeats.

### Changes in autism-related gene expression in brain regions of male mice with abnormal social behaviors induced by repeated agonistic interactions

Common changes in several gene expressions in the winners and losers, as well as gene expression changes specific for the different social groups and brain regions in mice have been identified in this experiment (Supplementary Table 1).

The *Tph2, Maoa, Slc6a4* and *Htr7* genes are associated with brain serotonergic activity and, according to several databases, are involved in numerous psychiatric disorders. Using the RT PCR method, we previously identified decreased expression of the *Tph2, Maoa*, and *Slc6a4* genes in the midbrain rapher nuclei of the winners [42] and losers [41]. The RNA-Seq transcriptome analysis data strongly confirmed these data for the *Tph2* and *Slc6a4* genes. We also identified decreased *Maoa* gene expression in the hypothalamus and increased expression of the *Tph2* and *Slc6a4* genes in the VTA of the winners and losers. *Htr7* gene expression was increased in the striatum of the losers.

It is well recognized that alterations in the peripheral and central indices of serotonin production, storage and signaling have associated with autism [64, 65]. Research evidence indicates that BDNF (brain-derived neurotrophic factor) promotes the survival and differentiation of serotonergic neurons [66]. Similar to these data, the involvement of BDNF in symptoms of ASDs in animal models [67-70] and autistic patients [71-73] has been demonstrated. In the current and previous studies [41-42], decreased *Bdnf* gene expression was identified in the raphe nuclei of the midbrain in the losers and winners. These findings provide confirmation of the supported hypothesis concerning the involvement of these genes in autistic spectrum behaviors. Changes in the expression of these genes in groups of mice with alternative social behaviors may regulate similar changes in the behaviors and emotional states, for example, reduced or low sociability, exploration deficits, reward dysfunction, and anxiety.

The *Shank* genes encode proteins that are members of the Shank family of synaptic proteins, which may function as molecular scaffolds in the postsynaptic density of excitatory synapses. Diseases associated with *Shank2* (*Auts17*) gene upregulation in the hippocampus of the losers include ASDs. Reduced *Shank3* expression is a major causative factor in several neurological symptoms. In the winners and losers, *Shank3* gene overactivation was identified in the hypothalamus, with reduced expression in the hippocampus of the winners, who demonstrate lower levels of social recognition and exploration, repetitive behaviors, and aggression. Thus, we suggest that the synthesis of Shank family synaptic proteins is disturbed in the hypothalamus and hippocampus in male mice with abnormal social behaviors.

The *Ube3a* gene encodes an E3 ubiquitin-protein ligase, which is a component of the ubiquitin protein degradation system. The expression of rodent *Ube3a* transcripts is associated with changes in the expression of the *Gabrb3* (*auts4*) gene, which encodes a protein that serves as the receptor for gamma-aminobutyric acid. We identified increased expression of the *Ube3a* gene in the midbrain of the winners and decreased expression of the *Ube3a* and *Gabrb3* (*auts4*) genes in the hypothalamus in both the winners and losers.

The *Reln* and *Pten* genes, as well as the *Ctnnd2, Nrxn1, Nrxn2, Nlgn1, Nlgn2*, and *Nlgn3* genes code proteins that are involved in focal or cell adhesion processes, which enable cells to interact and attach to a surface, substrate or another cell and are mediated by interactions between molecules of the cell surface. In both social groups, strongly decreased expressions of the *Reln* gene in the midbrain raphe nuclei and the *Pten* gene in the hypothalamus were identified. The winners exhibited decreased expression of the *Reln* gene in the hypothalamus, the *Pten* gene in the striatum, and the *Ctnnd2* and *Nrxn2* genes in the hippocampus, as well as increased expression of the *Nrxn1* gene in the midbrain raphe nuclei. The losers exhibited increased expression of the *Nrxn2, Nlgn2* and *Nlgn3* genes in the hypothalamus and the *Nlgn2* gene in the hippocampus, as well as decreased expression of the *Nrxn1* and *Nlgn1* genes in the hypothalamus, the *Reln* gene in the striatum and increased expression in the hippocampus. Increased expression of the *Pcdh10* gene, a potential calcium-dependent cell-adhesion protein, was identified in the hypothalamus and hippocampus of the losers and the *Cadps2* gene, which encodes a member of the calcium-dependent activator of secretion (CAPS) protein family, in the midbrain raphe nuclei of the winners and the VTA of the losers. It has been demonstrated that the *Bdnf* gene has pathways with the *Cadps2, Reln, Ctnnd2*, and *Nlgn1* genes.

Dysfunction of several genes is strongly suspected as a genetic cause of autism or ASDs. The *Mecp2* (*autsx3*) gene plays an essential role in mammalian development, and the MeCP2 protein binds to forms of DNA that have been methylated. The role of MECP2 in disease is primarily associated with a loss of function of the *Mecp2* gene or over expression. We identified increased expression of the *Mecp2* gene in the VTA of the losers and decreased expression in the hypothalamus in both the winners and losers.

The *Fmr1* gene, fragile X Mental Retardation Protein 1, encodes a protein that binds RNA and is associated with polysomes. Loss of function of the *Fmr1* gene as a result of microdeletion is responsible for the clinical phenotype of fragile X syndrome (FXS) [74]. In our study, hypoexpression of the *Fmr1* gene was identified in the hypothalamus of both social groups and activation of expression in the midbrain raphe nuclei of the winners. Animals in both social groups demonstrated cognitive dysfunctions, social anxiety, and stereotypic behaviors. The winners also exhibited aggression and hyperactivity, attention deficit behavior, obsessive behavior to attack conspecifics, and irritability. Similar to children with FXS, aggressive mice exhibit hypersensitivity to visual, auditory, tactile, and olfactory stimuli. The losers exhibited social withdrawal behaviors, including avoidance, indifference, and depression. Interestingly, a reduction in *Bdnf* expression in Fmr1 knockout mice worsens cognitive deficits and improves hyperactivity and sensorimotor deficits [70].

Decreased expression of the *Arx* gene in the hypothalamus of the winners and increased expression in the hippocampus and decreased expression in the striatum of the losers were identified in animals with abnormal social behaviors. It is well established that mental retardation is associated with mutations of the *Arx* gene, which has been described in several patients with autism or autistic features and causes a diverse spectrum of other diseases that includes cognitive impairment and epilepsy.

We identified downregulation of the *Foxp2* gene in the midbrain raphe nuclei of the winners and losers, in the hypothalamus of the winners and overexpression in the VTA of the losers. *Auts2* - Autism Susceptibility Candidate 2 gene, is thought to be a transcription factor required for normal brain development. It has been implicated in neurodevelopment and as a candidate gene for numerous neurological disorders that result from *Auts2* gene deficiency. *Auts2* gene expression was decreased in the hippocampus of the winners and increased in the hypothalamus of the losers.

*Ptchd1 (autsx4)* gene diseases, which are associated with PTCHD1 protein, include autism x-linked 4 and intellectual disability. There was decreased expression in the hypothalamus in both social groups. In the striatum decreased expression was specific only for the winners, with increased expression in the VTA of the losers. The *Oxtr* gene, which codes the oxytocin receptor, is significantly associated with ASD. Decreased expression of this gene was identified in the midbrain raphe nuclei of the winners and losers. The *En2* (*auts1*) gene is a Homeobox-containing gene and is thought to have a role in controlling development and contributes to as many as 40% of ASD cases, approximately twice the prevalence in the general population [75]. Increased expression of this gene was identified in the striatum and VTA.

Thus, the analysis of changes in expression of autism-related genes in different brain regions indicated interesting features. In the hippocampus, all autistic genes that changed their expression were not overlapping in male mice with alternative social behaviors. Moreover, all genes maintain decreased expression in the winners and increased expression in the losers. Previously, most of the various *Rps* and *Rpl* genes were upregulated in the winners [76] and downregulated in the losers [77] in this brain region and also did not overlap, with the exception of one *Rps39* gene. We associated this phenomenon with the activation of neurogenesis identified in the winners [78] and its decreased cell division and neurogenesis in the losers [79-82]. The hippocampus is responsible for cognitive function, memory consolidation and storage, and we suspect different cognitive processes in the winners and losers.

The VTA, which is widely implicated in the brain natural reward circuitry and emotions, exhibited increased expression of all genes in the animals with alternative social behaviors. Our previous behavioral data demonstrated reward dysfunction in both the winners and losers [34, 62]. In the striatum, which is responsible for the regulation of motor activity and stereotypical behaviors, we identified decreased expression of most genes that were not overlapping, which may be connected with the disturbances in psychomotor activity. We suspect responsibility of these genes for the stereotypic behaviors identified in both social groups. In the midbrain raphe nuclei, which comprise a multifunctional brain region that contains the majority of serotonergic neuronal bodies, we identified decreased expression of genes in the winners and losers and upregulation of specific genes in the winners only. In the hypothalamus, decreased gene expression was identified in the winners and losers, as well as upregulation of specific genes in the losers.

Autism is considered a multifactorial disorder that involves many genes which form the susceptibility to autism. Several of these genes changed their expression in the winners and losers under agonistic interactions, for example, *Auts1*; the *Auts17* gene, which is associated with a mutation in the *Shank2* gene; the *Autsx1* gene, which is associated with mutations in the *Nlgn3* gene; the *Autsx3 gene*, which is associated with mutations in the *Mecp2* gene; and *Autsx4*, which is associated with variation in the region of the chromosome that contains the *Ptchd1* gene. Considering all autism-related genes investigated, we suggest that the social environment may influence their expression, thereby forming symptoms similar with ASDs in adults. The combination of a neurochemical environment specific for every brain region with different functions may influence the specificity or similarity of changes in the behaviors of animals with alternative social behaviors.

### Alternative splicing expression profiling

Evidence indicates that alternative splicing contributes to ASDs [83, 84]. It has been suggested that alternative splicing characterized by aberrant mRNA expression of specific genes, such as the *Shank3, Nrxn1, Nlgn3, Nlgn4X* and other genes, may play a role in the ASDs [85-87].

In addition, a range of mutations located within splice sites or splicing factor binding sites of autism related genes [88] lead to abrupt splicing alterations; thus, we may assume that even in the absence of deleterious mutations, a stress-induced mis-regulation of alternatively spliced transcripts may influence the complex homeostasis of mRNA isoforms in brain tissues [89]. The highest emergence of isoforms in the 27 selected gene samples was identified in synapse specialized genes as previously reported and provides flexible neuronal communication [90]; thus, we assume that alterations in the splicing specific isoform expression profiles may lead to abnormal neurotransmitter behaviors that imply ASD associated genes, such as neurexins, neuroligins, *Shank1-3*, and *Pcdh10*. It should be noted that isoforms often coordinately express, particularly in synapse genes [89, 90]. Moreover, we identified the coordinated expression of *Nrxn2* and *Nlgn2* specific isoforms in the midbrain raphe nuclei and hypothalamus (Supplementary Fig. 1). Importantly, the ‘loser’ mice exhibited the most intense expression, which indicates that the signal of inhibitory synapses is increased [49] in these defeated mice. Thus, we underline the importance of paying attention to isoform co-expression profiling. These findings imply that autistm-related genes may be involved in the disturbances of social behaviors induced by chronic agonistic interactions in adult mice.

## Conclusion

As previously indicated, ASDs may comprise a substantially broader phenotype that comprises the less severe disorders than autism as developmental disorders, which includes individuals with symptoms of autism who do not meet the full criteria for autism or other disorders. Moreover, according to the APA, in the DSM-V, various affective and neurological disorders, such as depression, anxiety, schizophrenia, and ADHD, as well as many somatic disorders may be accompanied by impairments in social behaviors [2]. Based on an investigation of the networks of genes that represent their commonness or uniqueness for the impairment of social behaviors and bipolar disorder, authors [91] have hypothesized that comorbidity may be correlated with the genes involved in the gene pathways common to both disorders. It may be assumed that common genes may determine the disturbances in social behaviors similar to “autistic-like symptoms” under the psychopathology of aggressive behavior similar to psychosis, as demonstrated in the winners with chronic experience of aggression [34] or an anxiety and depression-like state in defeated mice, which develop under chronic negative social experiences in animals [30, 33, 35, 92]. Nevertheless, our behavioral approach mimicked many signs of impaired social behaviors in male mice, as well as changes in autism-associated gene expression, which may be useful for investigating brain molecular changes that may determine the subsequent psychoemotional state, cognitive dysfunction and disturbances in social behaviors. We suggest that abnormal social behaviors and changes in gene expression are a result of comorbidity with escalated aggression, depression- and anxiety-like state development as previously demonstrated [30, 33, 34, 92] in male mice under repeated agonistic interactions.

We assessed the RNA-Seq data consistency in the target genes using two approaches. First, we built up the Principal Components variation plot and identified brain region specific compact clustering, which thus confirmed the consistent expression pattern of the 45 samples (Fig. 9). Second, we visualized the Pearson pairwise correlation matrix via a Multidimensional Scaling (MDS) Plot (Fig. 11). We also identified segregation of the synapse genes and serotonergic genes, as well as other compact gene clusters, which supports functional co-variation of genes based on 45 observations, compared with the String gene association network (Fig. 1) based on external publications, databases, and experimental data. Thus, we assume that the RNA-Seq data adequately represents functioning of neurospecific genes in the 45 samples considered.

Changes in gene expression may also be caused in adults by different environments and may persist for a long time. Evidence in support of the fundamental role of this impact has been obtained in experiments on the molecular consequences of chronic agonistic interactions. The persistence of changed social behaviors [56] and changed expression of many genes have been demonstrated for the losers [41, 92] and winners [93]. Changes in the functional state of “specific autism genes” or genes of the neurotransmitter systems, which are involved in communication mechanisms and/or associated behavior patterns, may appear as a hereditary impairment and may explain the epigenetic nature of autism mechanisms discussed by researchers [7, 94-96]. These novel insights open broader perspectives for the experimental studies of the molecular mechanisms of ASDs and the identification of therapeutic approaches for pharmacological correction.

## Abbreviations

ASDs: autism spectrum disorders
VTA: ventral tegmental area
MRN: midbrain raphe nuclei
STR: striatum
HIP: hippocampus
HPT: hypothalamus
Los-10, Los-20: groups of chronically defeated mice after 10 or 20 days of agonistic interactions
Win-10, Win-20: groups of chronically victorious mice demonstrating daily aggression, during 10 or 20 days
FPKM: fragments per kilobase of transcript per million mapped reads
PC: Principal components
AHC: Agglomerative Hierarchical Clustering
MDS: Multidimentional Scaling

## Ethical Approval

All procedures were in compliance with the European Communities Council Directive 210/63/EU on September 22, 2010. The study was approved by Scientific Council N 9 of the Institute of Cytology and Genetics SD RAS of March, 24, 2010, N 613.

## Competing interests

The authors declare that they have no conflicts of interest.

## Funding

This work was funded by Russian Science Foundation, no. 17-15-00003.

## Author Contributions

I. L.K. substantially contributes to behavioral data acquisition and analysis of transcriptomic data; I.L.K., D.A.S., and A.G.G. contributes to acquisition of animals with alternative social behaviors and brain materials; V.N.B. implemented bioinformatics assessing of RNA-Seq data, analyzed and interpreted data, and wrote the main manuscript text. NNK have made substantial contributions to conception and design, analysis and interpretation of data, wrote the main manuscript text. All authors have been involved in drafting the manuscript, have given final approval of the version to be published, agree to be accountable for all aspects of the work in ensuring that questions related to the accuracy or integrity of any part of the work are appropriately investigated and resolved. N.N.K and V.N.B. equally contribute to the paper.

## Acknowledgements

We thank N.N. Lang for technical assistance and Dr. V.A. Naprimerov for organizational support on the behavioral experiment.

## Supplementary materials

**Supplementary Figure 1.**
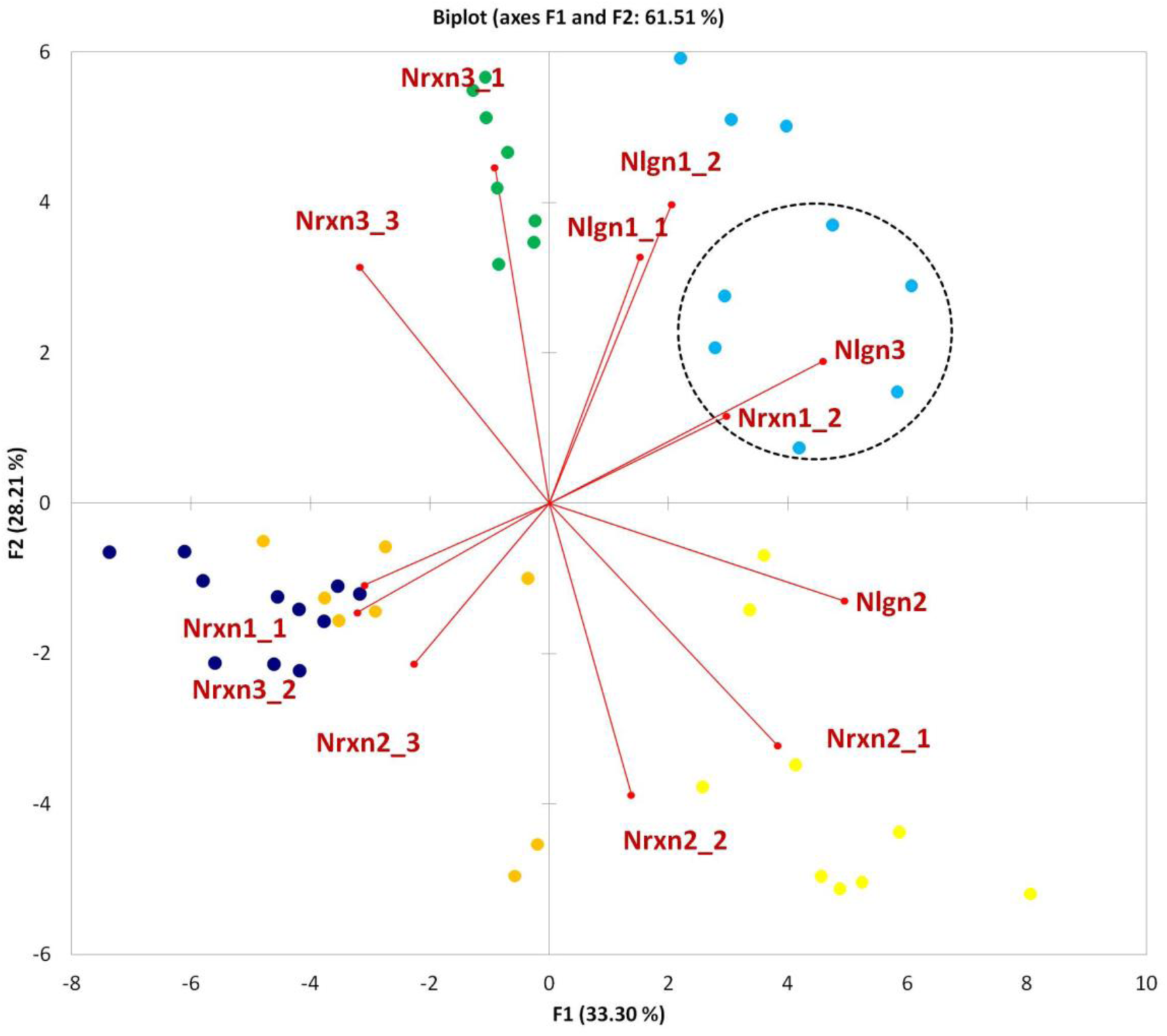
Illustration of Nrxn1,2,3 and Nlgn1,2 genes expression in brain regions based on RNA-Seq data for 45 mouse brain samples. The figure after dash in gene name denotes specific isoform. The Nlgn1 and Nrxn2 genes isoform pairs are concordant and located within each pair vicinity. In contrast, Nrxn1_1 is oppositely directed to its counterpart isoform Nrx1_2, manifestring brain region isoform specific expression. Brain region denotations: Green – hypothalamus; light blue – hippocampus; yellow – midbrain raphe nuclei; dark blue – ventral tegmental area (VTA); orange – striatum. Encircled are affected species in the hippocampus; they manifest increased Nrxn1_2 isoform expression. Control species don’t. Notably, alternative isoform Nrxn1_1 is attributable for Striatum/VTA. Nlgn2 is highly correlated with both isoforms of Nrxn2 and both manifest twofold higher expression than other genes considered in all tissues, but midbrain raphe nuclei: it manifests threefold higher expression of these genes. Isoforms encoding: (Nlgn1_1:NM_001163387; Nlgn1_2:NM_138666; Nlgn2:NM_198862; Nlgn3:NM_172932; Nrxn1_1:NM_020252; Nrxn1_2:NM_177284; Nrxn2_1:NM_001205234; Nrxn2_2:NM_001205235; Nrxn2_3:NM_020253; Nrxn3_1:NM_001198587; Nrxn3_2:NM_001252074; Nrxn3_3:NM_172544)

**Supplementary Table 1.** Differentially expressed autism-related genes in different brain regions in male mice with positive (winners) and negative (losers) social experience in daily agonistic interactions: RNAseq data

| Genes | Annotation | Winners | Losers | Skipping events |
| --- | --- | --- | --- | --- |
| *Tph2* | tryptophan hydroxylase 2 | ▼▼▼▼* MRN<br>▲▲▲▲* VTA | ▼▼▼▼* MRN<br>▲▲▲▲* VTA | 0 |
| *Maoa* | monoamine oxidase A | ▼ HPT | ▼▼▼▼* HPT | 1 |
| *Slc6a4* | solute carrier family 6 (serotonin transporter), member 4 | ▼▼▼▼* MRN<br>▲▲▲▲* VTA | ▼▼▼▼* MRN<br>▲▲▲▲* VTA | 0 |
| *Htr7* | 5-hydroxytryptamine (serotonin) receptor 7 |  | ▲▲▲▲* STR | 1 |
| *Shank2* | SH3 And Multiple Ankyrin Repeat Domains 2 (auts17) |  | ▲ HPC | 2 |
| *Shank3* | SH3/ankyrin domain gene 3 | ▲▲ HPT<br>▼▼ HPC | ▲▲▲* HPT | 3 |
| *Mecp2* | Methyl-CpG-Binding Protein 2 (autx3) | ▼ HPT | ▼ HPT<br>▲ VTA | 1 |
| *Ube3a* | ubiquitin protein ligase E3A | ▼ HPT<br>▲ MRN | ▼▼* HPT | 2 |
| *Gabrb3* | gamma-aminobutyric acid (GABA) A receptor, subunit beta 3 (auts4) | ▼ HPT | ▼ HPT | 0 |
| *Cntn3* | contactin 3 | ▲ MRN | ▼ HPT | 0 |
| *Reln* | reelin | ▼▼▼▼* MRN<br>▼▼ HPT | ▼▼▼▼* MRN<br>▲ HPC<br>▼ STR | 1 |
| *Pten* | phosphatase and tensin homolog | ▼▼* HPT<br>▼▼ STR | ▼▼▼* HPT | 1 |
| *Ctnnd2* | catenin (cadherin associated protein), delta 2 | ▼ HPC |  | 1 |
| *Nrxn1* | neurexin I | ▼ HPT<br>▲ MRN | ▼ HPT | 7 |
| *Nrxn2* | neurexin 2 | ▼▼ HPC | ▲▲▲* HPT | 11 |
| *Nlgn1* | neuroligin1 |  | ▼ HPT | 1 |
| *Nlgn2* | neuroligin2 |  | ▲ HPT<br>▲ HPC | 1 |
| *Nlgn3* | neuroligin3 (autsx1) |  | ▲ HPT | 3 |
| *Pcdh10* | protocadherin 10 |  | ▲ HPT<br>▲▲ HPC | 0 |
| *Cadps2* | Ca2+-dependent activator protein for secretion 2 | ▲ MRN | ▲▲ VTA | 3 |
| *Foxp2* | forkhead box P2 | ▼▼▼▼* MRN<br>▼ HPT | ▼▼▼▼* MRN<br>▲▲▲▲* VTA | 0 |
| *Fmr1* | fragile X mental retardation syndrome 1 homolog | ▼▼▼* HPT<br>▲ MRN | ▼▼* HPT | 1 |
| *Auts2* | autism susceptibility gene 2 protein | ▼ HPC | ▲▲▲* HPT | 3 |
| *En2* | engrailed 2 (auts1) |  | ▲▲ STR<br>▲▲▲▲* VTA | 0 |
| *Arx* | aristaless related homeobox | ▼▼ HPT | ▲▲▲▲* HPC<br>▼ STR | 0 |
| *Oxtr* | oxytocin receptor | ▼▼ MRN | ▼▼ MRN | 0 |
| *Ptchd1* | patched domain containing 1 | ▼▼▼▼* HPT<br>▼▼ STR | ▼▼▼▼* HPT<br>▲ VTA | 0 |

Note: ▼ - downregulation; ▲ - upregulation genes expression in comparison with the controls; ▲ $p \leq 0.05$ ; ▲▲ $p \leq 0.01$ ; ▲▲▲ $p \leq 0.001$ ; ▲▲▲▲ $p \leq 0.0001$ ; \* - $q < 0.05$ . MRN – midbrain raphe nuclei; HPC – hippocampus; STR – striatum; VTA – ventral tegmental area; HPT – hypothalamus. Gray colour – changes of genes expression common for both - the winners and losers.

**Supplementary Table 2.**
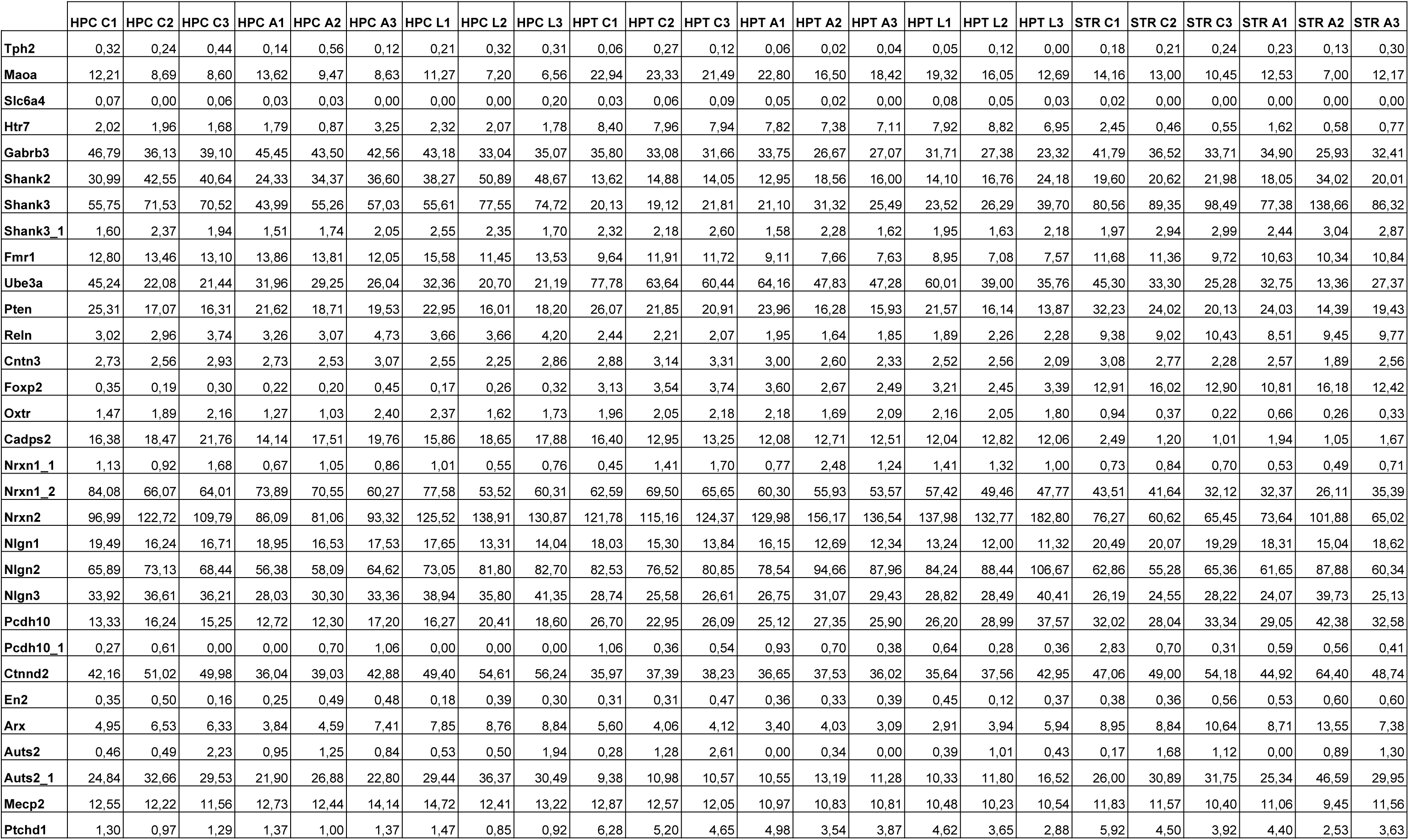
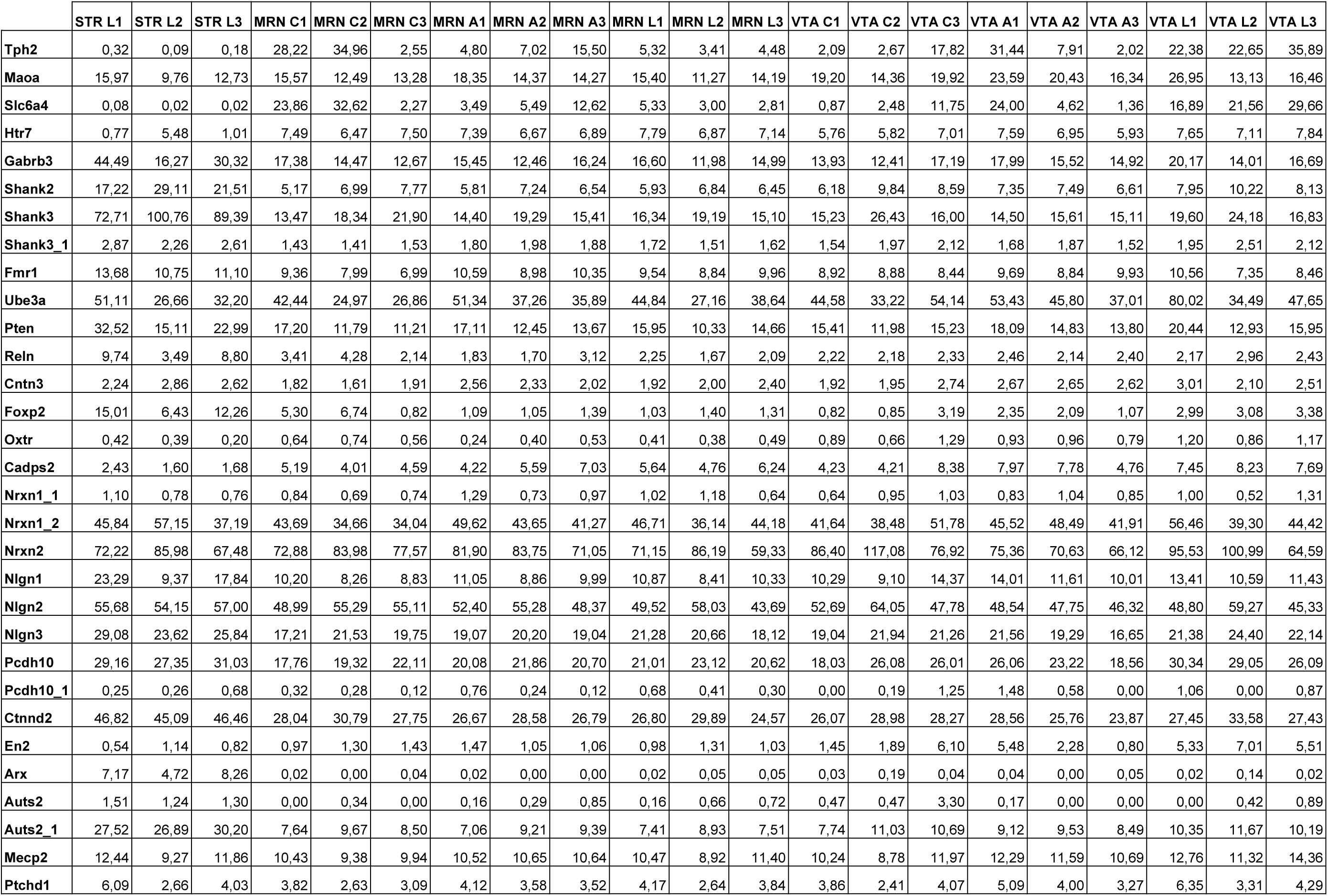
FPKM values for 30 transcripts analyzed in the study.

**Supplementary Table 3.** Pearson pairwise correlation matrix for the transcripts analyzed

| Variables | Tph2 | Maoa | Slc6a4 | Htr7 | Gabrb3 | Shank2 | Shank3 | Shank3 | Fmr1 | Ube3a | Pten | Reln | Cntn3 | Foxp2 | Oxtr | Cadps2 | Nrxn1_1 | Nrxn1_2 | Nrxn2 | Nlgn1 | Nlgn2 | Nlgn3 | Pcdh10 | Pcdh10_1 | Ctnnd2 | En2 | Arx | Auts2 | Auts2_1 | Mecp2 | Ptchd1 |
| --- | --- | --- | --- | --- | --- | --- | --- | --- | --- | --- | --- | --- | --- | --- | --- | --- | --- | --- | --- | --- | --- | --- | --- | --- | --- | --- | --- | --- | --- | --- | --- |
| **Tph2** | 1 | 0,285 | **0,993** | **0,407** | **-0,513** | **-0,470** | **-0,444** | -0,257 | **-0,349** | 0,180 | **-0,305** | -0,211 | -0,286 | -0,084 | -0,209 | -0,206 | -0,080 | -0,262 | **-0,326** | **-0,392** | **-0,492** | **-0,454** | -0,030 | 0,140 | **-0,495** | **0,743** | **-0,562** | -0,131 | **-0,429** | 0,049 | 0,205 |
| **Maoa** | 0,285 | 1 | 0,240 | **0,680** | -0,252 | **-0,681** | **-0,665** | -0,217 | **-0,338** | **0,930** | 0,182 | **-0,400** | 0,285 | -0,123 | 0,172 | -0,093 | 0,239 | 0,104 | 0,018 | -0,107 | -0,121 | **-0,493** | 0,134 | **0,333** | **-0,612** | **0,365** | **-0,525** | -0,177 | **-0,700** | 0,065 | **0,778** |
| **Slc6a4** | **0,993** | 0,240 | 1 | **0,400** | **-0,507** | **-0,460** | **-0,432** | -0,254 | **-0,359** | 0,141 | **-0,315** | -0,198 | **-0,328** | -0,072 | -0,211 | -0,208 | -0,090 | -0,276 | **-0,303** | **-0,406** | **-0,463** | **-0,435** | -0,033 | 0,102 | **-0,475** | **0,707** | **-0,548** | -0,155 | **-0,421** | 0,016 | 0,179 |
| **Htr7** | **0,407** | **0,680** | **0,400** | 1 | **-0,658** | **-0,697** | **-0,857** | **-0,499** | **-0,687** | **0,595** | **-0,407** | **-0,764** | -0,085 | **-0,475** | 0,177 | -0,004 | 0,286 | -0,035 | 0,200 | **-0,698** | -0,024 | **-0,506** | 0,018 | 0,074 | **-0,777** | **0,366** | **-0,777** | **-0,297** | **-0,908** | -0,254 | **0,423** |
| **Gabrb3** | **-0,513** | -0,252 | **-0,507** | **-0,658** | 1 | **0,686** | **0,536** | **0,337** | **0,731** | -0,043 | **0,770** | **0,436** | **0,520** | 0,220 | **0,463** | **0,494** | 0,066 | **0,579** | 0,221 | **0,890** | **0,394** | **0,680** | -0,134 | 0,144 | **0,685** | **-0,504** | **0,738** | 0,266 | **0,642** | **0,582** | -0,185 |
| **Shank2** | **-0,470** | **-0,681** | **-0,460** | **-0,697** | **0,686** | 1 | **0,710** | **0,326** | **0,650** | **-0,520** | 0,219 | 0,284 | 0,275 | 0,002 | **0,400** | **0,577** | -0,041 | **0,451** | **0,393** | **0,472** | **0,480** | **0,871** | -0,160 | -0,148 | **0,841** | **-0,415** | **0,761** | **0,300** | **0,836** | **0,399** | **-0,669** |
| **Shank3** | **-0,444** | **-0,665** | **-0,432** | **-0,857** | **0,536** | **0,710** | 1 | **0,660** | **0,481** | **-0,569** | **0,343** | **0,800** | 0,119 | **0,641** | -0,176 | -0,113 | -0,265 | -0,091 | -0,047 | **0,587** | 0,233 | **0,574** | **0,337** | 0,023 | **0,911** | **-0,361** | **0,896** | 0,293 | **0,959** | 0,065 | -0,290 |
| **Shank3** | -0,257 | -0,217 | -0,254 | **-0,499** | **0,337** | **0,326** | **0,660** | 1 | 0,247 | -0,137 | **0,346** | **0,671** | 0,146 | **0,672** | -0,123 | -0,203 | -0,056 | -0,171 | 0,006 | **0,525** | 0,233 | **0,361** | **0,588** | 0,045 | **0,651** | -0,042 | **0,672** | **0,369** | **0,622** | 0,186 | 0,145 |
| **Fmr1** | **-0,349** | **-0,338** | **-0,359** | **-0,687** | **0,731** | **0,650** | **0,481** | 0,247 | 1 | -0,183 | **0,545** | **0,317** | **0,437** | 0,041 | 0,177 | **0,365** | -0,043 | **0,574** | -0,076 | **0,643** | -0,035 | **0,461** | **-0,411** | -0,036 | **0,521** | **-0,361** | **0,511** | **0,334** | **0,613** | **0,593** | **-0,344** |
| **Ube3a** | 0,180 | **0,930** | 0,141 | **0,595** | -0,043 | **-0,520** | **-0,569** | -0,137 | -0,183 | 1 | **0,384** | **-0,360** | **0,401** | -0,114 | 0,283 | 0,052 | 0,277 | 0,283 | 0,117 | 0,057 | -0,008 | **-0,314** | 0,112 | **0,408** | **-0,474** | 0,273 | **-0,379** | -0,132 | **-0,588** | 0,222 | **0,766** |
| **Pten** | **-0,305** | 0,182 | **-0,315** | **-0,407** | **0,770** | 0,219 | **0,343** | **0,346** | **0,545** | **0,384** | 1 | **0,507** | **0,545** | **0,457** | 0,194 | 0,064 | -0,025 | **0,343** | -0,045 | **0,865** | 0,111 | 0,255 | 0,110 | **0,448** | **0,398** | -0,277 | **0,521** | 0,104 | **0,347** | **0,509** | **0,401** |
| **Reln** | -0,211 | **-0,400** | -0,198 | **-0,764** | **0,436** | 0,284 | **0,800** | **0,671** | **0,317** | **-0,360** | **0,507** | 1 | 0,005 | **0,892** | **-0,408** | **-0,452** | **-0,351** | **-0,403** | **-0,376** | **0,663** | -0,037 | 0,209 | **0,470** | 0,188 | **0,666** | -0,232 | **0,737** | 0,220 | **0,715** | 0,047 | 0,122 |
| **Cntn3** | -0,286 | 0,285 | **-0,328** | -0,085 | **0,520** | 0,275 | 0,119 | 0,146 | **0,437** | **0,401** | **0,545** | 0,005 | 1 | -0,027 | **0,500** | **0,399** | 0,262 | **0,591** | 0,121 | **0,500** | 0,135 | 0,205 | -0,053 | **0,413** | 0,207 | -0,058 | 0,269 | **0,344** | 0,137 | **0,580** | 0,183 |
| **Foxp2** | -0,084 | -0,123 | -0,072 | **-0,475** | 0,220 | 0,002 | **0,641** | **0,672** | 0,041 | -0,114 | **0,457** | **0,892** | -0,027 | 1 | **-0,459** | **-0,637** | -0,242 | **-0,513** | **-0,331** | **0,498** | -0,005 | 0,029 | **0,694** | 0,278 | **0,495** | -0,118 | **0,572** | 0,179 | **0,492** | -0,146 | **0,412** |
| **Oxtr** | -0,209 | 0,172 | -0,211 | 0,177 | **0,463** | **0,400** | -0,176 | -0,123 | 0,177 | 0,283 | 0,194 | **-0,408** | **0,500** | **-0,459** | 1 | **0,854** | **0,396** | **0,715** | **0,741** | 0,180 | **0,631** | **0,559** | -0,242 | 0,036 | 0,164 | -0,182 | 0,144 | 0,108 | -0,029 | **0,480** | -0,170 |
| **Cadps2** | -0,206 | -0,093 | -0,208 | -0,004 | **0,494** | **0,577** | -0,113 | -0,203 | **0,365** | 0,052 | 0,064 | **-0,452** | **0,399** | **-0,637** | **0,854** | 1 | **0,316** | **0,801** | **0,617** | 0,166 | **0,483** | **0,612** | **-0,520** | -0,118 | 0,189 | -0,194 | 0,127 | 0,150 | 0,096 | **0,542** | **-0,478** |
| **Nrxn1_1** | -0,080 | 0,239 | -0,090 | 0,286 | 0,066 | -0,041 | -0,265 | -0,056 | -0,043 | 0,277 | -0,025 | **-0,351** | 0,262 | -0,242 | **0,396** | **0,316** | 1 | **0,354** | **0,372** | -0,069 | **0,294** | 0,107 | -0,091 | 0,018 | -0,123 | -0,086 | -0,158 | 0,219 | -0,230 | 0,042 | 0,042 |
| **Nrxn1_2** | -0,262 | 0,104 | -0,276 | -0,035 | **0,579** | **0,451** | -0,091 | -0,171 | **0,574** | 0,283 | **0,343** | **-0,403** | **0,591** | **-0,513** | **0,715** | **0,801** | **0,354** | 1 | **0,424** | **0,316** | 0,277 | **0,446** | **-0,563** | -0,033 | 0,117 | -0,214 | 0,081 | 0,181 | 0,074 | **0,588** | -0,293 |
| **Nrxn2** | **-0,326** | 0,018 | **-0,303** | 0,200 | 0,221 | **0,393** | -0,047 | 0,006 | -0,076 | 0,117 | -0,045 | **-0,376** | 0,121 | **-0,331** | **0,741** | **0,617** | **0,372** | **0,424** | 1 | -0,066 | **0,902** | **0,655** | 0,099 | -0,139 | 0,276 | -0,265 | 0,206 | -0,049 | 0,040 | 0,025 | -0,216 |
| **Nlgn1** | **-0,392** | -0,107 | **-0,406** | **-0,698** | **0,890** | **0,472** | **0,587** | **0,525** | **0,643** | 0,057 | **0,865** | **0,663** | **0,500** | **0,498** | 0,180 | 0,166 | -0,069 | **0,316** | -0,066 | 1 | 0,162 | **0,470** | 0,107 | **0,312** | **0,634** | -0,285 | **0,726** | **0,300** | **0,635** | **0,553** | 0,101 |
| **Nlgn2** | **-0,492** | -0,121 | **-0,463** | -0,024 | **0,394** | **0,480** | 0,233 | 0,233 | -0,035 | -0,008 | 0,111 | -0,037 | 0,135 | -0,005 | **0,631** | **0,483** | **0,294** | 0,277 | **0,902** | 0,162 | 1 | **0,758** | **0,304** | -0,067 | **0,530** | **-0,459** | **0,511** | 0,033 | 0,275 | -0,006 | -0,162 |
| **Nlgn3** | **-0,454** | **-0,493** | **-0,435** | **-0,506** | **0,680** | **0,871** | **0,574** | **0,361** | **0,461** | **-0,314** | 0,255 | 0,209 | 0,205 | 0,029 | **0,559** | **0,612** | 0,107 | **0,446** | **0,655** | **0,470** | **0,758** | 1 | 0,066 | -0,095 | **0,810** | **-0,395** | **0,747** | 0,233 | **0,696** | **0,363** | **-0,492** |
| **Pcdh10** | -0,030 | 0,134 | -0,033 | 0,018 | -0,134 | -0,160 | **0,337** | **0,588** | **-0,411** | 0,112 | 0,110 | **0,470** | -0,053 | **0,694** | -0,242 | **-0,520** | -0,091 | **-0,563** | 0,099 | 0,107 | **0,304** | 0,066 | 1 | **0,312** | 0,283 | 0,162 | **0,336** | 0,028 | 0,172 | -0,281 | **0,520** |
| **Pcdh10_1** | 0,140 | **0,333** | 0,102 | 0,074 | 0,144 | -0,148 | 0,023 | 0,045 | -0,036 | **0,408** | **0,448** | 0,188 | **0,413** | 0,278 | 0,036 | -0,118 | 0,018 | -0,033 | -0,139 | **0,312** | -0,067 | -0,095 | **0,312** | 1 | -0,021 | 0,215 | 0,091 | -0,069 | -0,041 | 0,204 | **0,502** |
| **Ctnnd2** | **-0,495** | **-0,612** | **-0,475** | **-0,777** | **0,685** | **0,841** | **0,911** | **0,651** | **0,521** | **-0,474** | **0,398** | **0,666** | 0,207 | **0,495** | 0,164 | 0,189 | -0,123 | 0,117 | 0,276 | **0,634** | **0,530** | **0,810** | 0,283 | -0,021 | 1 | **-0,444** | **0,961** | **0,335** | **0,937** | 0,210 | **-0,334** |
| **En2** | **0,743** | **0,365** | **0,707** | **0,366** | **-0,504** | **-0,415** | **-0,361** | -0,042 | **-0,361** | 0,273 | -0,277 | -0,232 | -0,058 | -0,118 | -0,182 | -0,194 | -0,086 | -0,214 | -0,265 | -0,285 | **-0,459** | **-0,395** | 0,162 | 0,215 | **-0,444** | 1 | **-0,511** | 0,031 | **-0,360** | 0,123 | 0,259 |
| **Arx** | **-0,562** | **-0,525** | **-0,548** | **-0,777** | **0,738** | **0,761** | **0,896** | **0,672** | **0,511** | **-0,379** | **0,521** | **0,737** | 0,269 | **0,572** | 0,144 | 0,127 | -0,158 | 0,081 | 0,206 | **0,726** | **0,511** | **0,747** | **0,336** | 0,091 | **0,961** | **-0,511** | 1 | 0,281 | **0,901** | 0,253 | -0,201 |
| **Auts2** | -0,131 | -0,177 | -0,155 | **-0,297** | 0,266 | **0,300** | 0,293 | **0,369** | **0,334** | -0,132 | 0,104 | 0,220 | **0,344** | 0,179 | 0,108 | 0,150 | 0,219 | 0,181 | -0,049 | **0,300** | 0,033 | 0,233 | 0,028 | -0,069 | **0,335** | 0,031 | 0,281 | 1 | **0,320** | 0,224 | -0,136 |
| **Auts2_1** | **-0,429** | **-0,700** | **-0,421** | **-0,908** | **0,642** | **0,836** | **0,959** | **0,622** | **0,613** | **-0,588** | **0,347** | **0,715** | 0,137 | **0,492** | -0,029 | 0,096 | -0,230 | 0,074 | 0,040 | **0,635** | 0,275 | **0,696** | 0,172 | -0,041 | **0,937** | **-0,360** | **0,901** | **0,320** | 1 | 0,233 | **-0,431** |
| **Mecp2** | 0,049 | 0,065 | 0,016 | -0,254 | **0,582** | **0,399** | 0,065 | 0,186 | **0,593** | 0,222 | **0,509** | 0,047 | **0,580** | -0,146 | **0,480** | **0,542** | 0,042 | **0,588** | 0,025 | **0,553** | -0,006 | **0,363** | -0,281 | 0,204 | 0,210 | 0,123 | 0,253 | 0,224 | 0,233 | 1 | -0,047 |
| **Ptchd1** | 0,205 | **0,778** | 0,179 | **0,423** | -0,185 | **-0,669** | -0,290 | 0,145 | **-0,344** | **0,766** | **0,401** | 0,122 | 0,183 | **0,412** | -0,170 | **-0,478** | 0,042 | -0,293 | -0,216 | 0,101 | -0,162 | **-0,492** | **0,520** | **0,502** | **-0,334** | 0,259 | -0,201 | -0,136 | **-0,431** | -0,047 | 1 |

